# Genome-Wide Investigation of Transcription Factor Occupancy and Dynamics Using cFOOT-seq

**DOI:** 10.1101/2025.07.17.664523

**Authors:** Heng Wang, Ang Wu, Meng-Chen Yang, Di Zhou, Xiyang Chen, Zhifei Shi, Yiqun Zhang, Yu-Xin Liu, Kai Chen, Xiaosong Wang, Xiao-Fang Cheng, Baodan He, Yutao Fu, Lan Kang, Yujun Hou, Kun Chen, Shan Bian, Juan Tang, Jianhuang Xue, Chenfei Wang, Xiaoyu Liu, Jiejun Shi, Shaorong Gao, Jia-Min Zhang

## Abstract

Gene regulation relies on the precise binding of transcription factors (TFs) at regulatory elements, but simultaneously detecting hundreds of TFs on chromatin is challenging. We developed cFOOT-seq, a cytosine deaminase-based TF footprinting assay, for high-resolution, quantitative genome-wide assessment of TF binding in both open and closed chromatin, even with small cell numbers. By utilizing the dsDNA deaminase SsdA_tox_, cFOOT-seq converts accessible cytosines to uracil while preserving genomic integrity, making it compatible with techniques like ATAC-seq for sensitive and cost-effective detection of TF occupancy at single-molecule and single-cell level. Our approach enables the delineation of TF footprints, quantification of occupancy, and examination of chromatin influences. Notably, cFOOT-seq, combined with FootTrack analysis, predicts TF occupancy dynamics. We demonstrate its application in capturing cell type-specific TFs, analyzing TF dynamics during reprogramming, and revealing TF dependencies on chromatin remodelers. Overall, cFOOT-seq represents a robust approach for investigating the genome-wide dynamics of TF occupancy and elucidating the cis-regulatory architecture underlying gene regulation.

## Introduction

Chromatin accessibility, nucleosome arrangement, and transcription factors binding to cis-regulatory elements shape the genome’s regulatory landscape and dictate transcriptional activity^1-3^. Mapping the genomic binding sites of all active transcription factors and their dynamic changes during cellular processes and environmental responses is crucial for understanding their roles in cell identity and the reshaping of gene regulatory networks^4-7^. However, achieving this comprehensive mapping remains challenging due to the lack of sensitive and robust methods for large-scale dynamic assessment of genomic TF binding.

DNA binding specificities of TF can be deduced by in vitro methods quantifying TF-DNA interaction by sequencing such as SELEX^8, 9^, however, it can’t provide the in vivo localization information of TFs. Occupancy of specific transcription factors on chromatin in vivo can be profiled by ChIP-seq^10, 11^, or its optimized strategies such as ChIP-exo^12, 13^. Recently developed enzyme-tethering and cutting or tagging-dependent methods, including CUT&RUN^14^, ChIL-seq^15^, CUT&TAG^16^, ACT-seq^17^ and CoBATCH^18^ provide genome-wide distributions of chromatin-binding proteins with improved signal-to-noise ratios and lower sample requirements. Recent DNA-modifying enzyme-based approaches, such as DiMeLo-seq^19^, nanoHiMe-seq^20^, and BIND&MODIFY^21^, combine antibody-based protein recognition with protein A-fused non-specific DNA adenine methyltransferases to map histone modifications and protein-DNA interactions, with methylation detection performed using PacBio or Nanopore technologies. While improvement on the throughput for single-cell and multiple targets^22-29^, the antibody-based strategies are still constrained by the necessity for highly specific antibodies and challenges in scalability, which make them hard to study the genomic kinetics of TF binding events for hundreds of TFs simultaneously.

TF binding can also be inferred from chromatin footprints caused by TF occupancy^30, 31^. DNase-seq^32-35^ and ATAC-seq^36-39^ have been used to detect the footprints of TF by identifying regions protected by TF from nuclease cleavage. Specifically, DNase-seq has been utilized in the ENCODE project to detect human TF footprints across hundreds of cell types^40^. However, DNase-seq and ATAC-seq are dependent on numerous cutting events on open chromatin to robustly examine the protection effect from TF, which make it hard to robustly detect TF occupancy on less accessible chromatin and in lower number of cells, because of relative sparse cutting events. More recent advancements, including Single Molecule Footprinting (SMF)^41, 42^, SMAC-seq^43^, and Fiber-seq^44^, employ DNA methyltransferases to modify accessible cytosines or adenines without DNA cleavage. However, SMF’s utility is limited by the sporadic occurrence of CpG and GpC sites, interference from endogenous cytosine methylation, and DNA degradation due to bisulfite conversion^31^. SMAC-seq and Fiber-seq address these issues by employing adenine methyltransferase and detecting DNA modifications via nanopore or PacBio sequencing^43, 44^. Despite these improvements, third-generation sequencing methods are still hindered by accuracy, throughput, and cost, and require large cell quantities due to the inability to amplify modified DNA prior to sequencing.

Here, we present cFOOT-seq, a cytosine deaminase-based footprinting assay for occupancy and organization of transcription factors by sequencing (cFOOT-seq), that simultaneously assesses chromatin accessibility, nucleosome positioning, and the occupancy of hundreds of transcription factors (TFs). cFOOT-seq leverages the dsDNA cytosine deaminase SsdA_tox_ to convert accessible cytosines to uracil, encoding chromatin organization directly into DNA sequences. The positions of nucleosomes and TFs are inferred based on their protective effects against DNA deamination. cFOOT-seq is highly compatible with ATAC-seq, enabling cost-effective detection of TF occupancy in open chromatin and supporting detection of TF binding at single-molecule and single-cell level.

cFOOT-seq not only examines the dynamics of TF occupancy at known sites, but also provides de novo prediction of TF binding sites with FootTrack analysis of TF footprints and motifs. With cFOOT-seq, we analyzed the impact of chromatin context on TF occupancy, and detected the dynamics of TF binding in the early stages of OSKM-mediated reprogramming of mouse embryonic fibroblasts (MEFs). We further defined the dependence of more than one hundred TFs on the SWI/SNF chromatin remodeling complex in HepG2, and revealed an observation of spatial organization that TFs with the similar SWI/SNF dependency are frequently located in close proximity on the chromatin. We anticipate that future applications of cFOOT-seq will provide new insights into decoding the grammar of TF binding on chromatin and constructing accurate gene regulatory networks.

### Design

cFOOT-seq was developed to address the need for a sensitive and scalable method to map TF binding, chromatin accessibility, and nucleosome positioning^1, 31^. Traditional methods like ChIP-seq^11^ and CUT&Tag^16^ rely on antibodies, limiting scalability, while nuclease-based approaches like DNase-seq^32^ and ATAC-seq^38^ struggle with detecting TF binding in less accessible chromatin and require large cell inputs. Additionally, single-molecule and single-cell analyses remain challenging with these bulk-sequencing-dependent techniques^45^.

To address these limitations, cFOOT-seq leverages the dsDNA cytosine deaminase SsdA_tox_, which converts accessible cytosines to uracils, encoding chromatin structure and TF occupancy directly into DNA sequence changes. This design eliminates the need for nuclease cleavage or antibodies, preserving DNA integrity while enabling the detection of TF footprints in both open and closed chromatin regions. cFOOT-seq is highly adaptable and can be integrated with complementary approaches, such as ATAC-seq, to enrich for open chromatin regions. This integration enhances detection sensitivity and reduces sequencing costs. Notably, the combined ATAC-cFOOT workflow extends the utility of cFOOT-seq to single-molecule and single-cell level, enabling detailed mapping of TF binding at unprecedented resolution. cFOOT-seq can examines the dynamics of TF occupancy at known sites, and provides de novo prediction of TF binding sites with FootTrack analysis. By addressing limitations related to cell number, chromatin openness, and enzyme biases, cFOOT-seq provides a more accurate and comprehensive understanding of TF dynamics and chromatin architecture across the genome.

## Result

### cFOOT-seq maps chromatin accessibility, nucleosome positioning and TF footprint

To convert chromatin architecture into DNA sequences at single-nucleotide resolution, we developed cFOOT-seq, a cytosine deaminase-based genomic footprinting assay that depicts chromatin accessibility, nucleosome positioning, and transcription factor occupancy via sequencing. In cFOOT-seq, permeabilized cells are treated with double-stranded DNA (dsDNA) cytosine deaminases, which preferentially convert accessible cytosine (C) to uracil (U). PCR amplification subsequently results in C-to-T conversions. Open chromatin, being more susceptible to deamination, shows higher conversion rate, indicating chromatin accessibility. Conversely, nucleosome and TF binding protect the DNA from deamination, revealing their footprints as regions with decreased conversion rate. (Fig. 1A).

**Figure 1:**
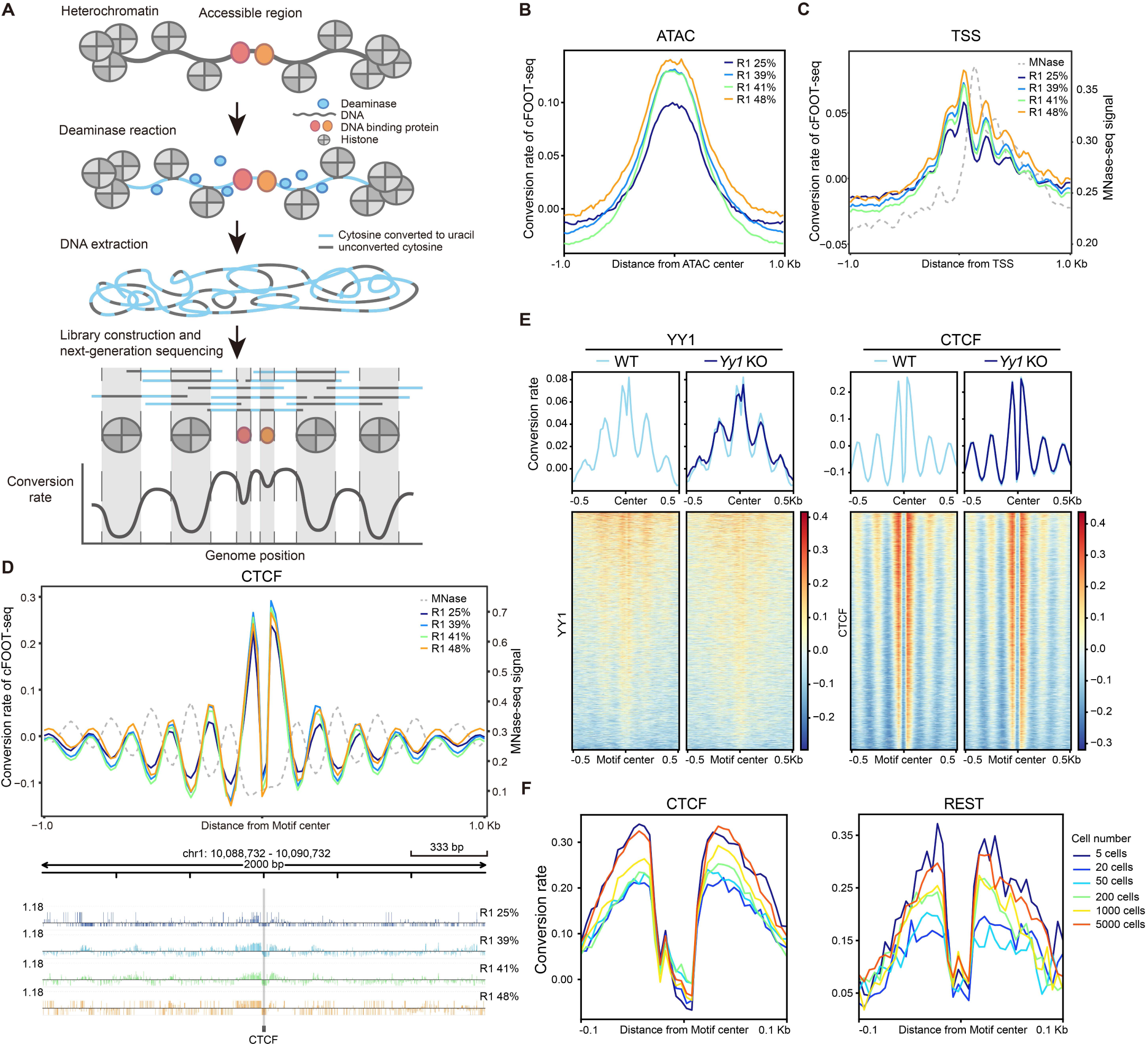
cFOOT-seq maps chromatin accessibility, nucleosome occupancy, and TF footprints. **A.** Schematic of the cFOOT-seq workflow. After cell permeabilization, dsDNA deaminases (blue) convert cytosine to uracil in accessible chromatin DNA (blue lines), but DNA occupied by nucleosomes or transcription factors (TFs) (gray lines) remains unconverted. The profile of DNA conversion rate defines nucleosome occupancy and TF footprints. **B.** Average profile of normalized DNA conversion rate around the centers of open chromatin regions (OCRs) which are defined by ATAC-seq peaks in R1 cells, showing samples with increasing average genomic conversion rate. **C.** Average profile of normalized DNA conversion rate of cFOOT-seq and MNase-seq signal around transcription start sites (TSS) in R1 cells, showing samples treated with increasing average genomic conversion rate. The pattern of nucleosome positioning, as demonstrated by MNase-seq, is depicted by the grey dashed line. **D.** Average profiles of normalized DNA conversion rate of cFOOT-seq and MNase-seq signal around CTCF binding sites defined by ChIP-seq in R1 cells, displaying the nucleosome position and CTCF footprint. The pattern of nucleosome positioning as indicated by MNase-seq is shown with a grey dashed line (top). Integrative Genomic Viewer (IGV) browser view of a representative site (chr1:10,088,732-10,090,732) flanking the CTCF motif showing the distribution of DNA conversion rate around CTCF motifs at single site (bottom). The position of CTCF motif is marked by gray bar. **E.** Average profiles of normalized DNA conversion rate around YY1 (left) and CTCF (right) binding motifs in wild-type (WT, 33%) and *Yy1* knockout (KO, 32%) R1 cells, displaying the changes of YY1 footprint after knockout of *Yy1*. Heatmaps showing DNA conversion rate around individual binding sites aligned to motif centers and ranked by average conversion rate within ±0.1 kb of the motif center (bottom). The binding sites of YY1 and CTCF are defined by ChIP-seq in R1. **F.** Average profiles of normalized DNA conversion rate around CTCF (left) or REST (right) binding motifs in R1 samples with low input cell numbers (5-5000), displaying the footprints of CTCF and REST at their known binding sites. The binding sites of CTCF and REST are defined by ChIP-seq in R1. See also Figures S1, S2

For mapping the chromatin landscape and TF footprints across the genome, DNA deaminases with robust enzymatic activity and minimal sequence bias on dsDNA are essential. While DddA_tox_ was initially identified for its ability to deaminate cytosine in dsDNA, its TC context preference limits its use for TF footprint detection^46^. SsdA_tox_^47^, along with recently identified DddA_tox_ homologs like Ddd_Ss^48^, offer more potent enzymatic activity and reduced sequence bias. To evaluate their activity and sequence bias for cFOOT-seq, we purified SsdA_tox_, Ddd_Ss, and another DddA_tox_ homolog, Ddd_Fa, for comparison (Figure S1A)

Our findings demonstrate that SsdA_tox_ exhibits more consistent deamination across a range of DNA oligonucleotides compared to Ddd_Ss and Ddd_Fa (Figure S1B), with significantly lower sequence bias. In tests using naked genomic DNA from R1 cells, SsdA_tox_ shows rapid deamination, achieving near 100% conversion at a relatively low enzyme concentration, while Ddd_Ss and Ddd_Fa exhibit a more gradual increase in conversion rate with increasing enzyme concentration (Figure S1C-D). This highlights SsdA_tox_’s superior catalytic efficiency compared to Ddd_Ss and Ddd_Fa.

Nucleotide preference studies reveal that Ddd_Fa retains a strong TC context bias, while Ddd_Ss exhibits less sequence specificity (Figure S1E and Figure S1F). In contrast, SsdA_tox_ shows minimal sequence bias, making it ideal for comprehensive genomic analysis (Figure S1F). When assessing CTCF motifs on R1 naked genomic DNA, SsdA_tox_ demonstrated consistent performance with minimal sequence bias, whereas Ddd_Ss and Ddd_Fa showed greater variability, reinforcing SsdA_tox_’s suitability for precise TF footprint detection (Figure S1G).

Moreover, SsdA_tox_ is less affected by DNA methylation status compared to Ddd_Ss and Ddd_Fa, as evidenced by its deamination efficiency at both high and low methylation sites (Figure S1H). Mass spectrometry analysis of deamination at C and 5mC sites further supports that SsdA_tox_ deaminates 5mC more efficiently than the other enzymes (Figure S1I). This characteristic is crucial for detecting TF footprints in DNA regions with prevalent methylation. With its high activity and minimal bias, SsdA_tox_ is the optimal choice for use in cFOOT-seq, ensuring accurate representation of chromatin organization across broad sequence coverage.

Treating permeabilized R1 cells with increasing SsdA_tox_ concentrations, we observed higher genomic conversion rate correlating with higher SsdA_tox_ concentrations (Fig. S2A). Conversion rate at ATAC-seq peaks and transcription start sites (TSS) were higher than those in flanking regions (Fig. 1B-C), suggesting that these conversion rate reflect chromatin accessibility. This pattern was consistent across multiple cell types, including human HepG2 cells, further demonstrating the robustness of the assay (Fig. S2B-C)

cFOOT-seq detected nucleosome phasing patterns around TSS and CTCF binding sites, consistent with the nucleosome positioning signal from MNase-seq (Fig. 1C-D). Conversion rate at TSS revealed downstream nucleosome positioning patterns (Fig. 1C). The conversion rate around CTCF motif also showed strong nucleosome phasing patterns, consistent across samples with different conversion rate after normalization (Fig. 1D). Similar nucleosome phasing patterns were observed around the binding sites of REST and YY1 (Fig. S2D-E), indicating a unique chromatin landscape around their binding sites.

cFOOT-seq was able to detect TF occupancy sites based on their footprints. Conversion rate at the CTCF motif were significantly lower than those around the motifs, representing CTCF footprints at binding sites due to protection from deamination (Fig. 1A, 1D). Footprints were also detected at REST and YY1 binding sites (Fig. S2D-E). Knocking out YY1 in R1 cells led to a notable reduction in both nucleosome phasing and footprint signals around YY1 binding motifs, while signals around CTCF motifs were unaffected, confirming YY1 occupancy signals detected by cFOOT-seq are specific to YY1 (Fig. 1E, Fig. S2F). Moreover, the loss of YY1 footprints can also be observed at individual binding sites in Yy1 KO R1 cells, further supporting the specificity of the detected footprint signals (Fig. S2G).

cFOOT-seq captures both nucleosome positions and TF footprints through deamination, rather than cutting chromatin DNA, providing a more comprehensive view of chromatin status in both open chromatin regions (OCR) and closed chromatin regions (CCR). Compared to DNase-seq and ATAC-seq, cFOOT-seq successfully detects CTCF footprints in both OCR and CCR, revealing nucleosome phasing around the CTCF binding sites (Fig. S2H). While ATAC-seq and DNase-seq can detect CTCF footprints in OCR, their ability to identify footprints in CCR is limited, and they lack detailed chromatin context, such as nucleosome phasing. Similarly, cFOOT-seq detects TF CEBPA footprints in both chromatin contexts, further highlighting its advantage in simultaneously capturing chromatin structure and TF occupancy across the genome over DNase-seq and ATAC-seq.

Finally, we assessed cFOOT-seq’s ability to characterize chromatin landscapes using limited cell numbers. Remarkably, the assay could detect chromatin accessibility, nucleosome phasing patterns around CTCF, and footprints of CTCF and REST in as few as 5 to 5000 R1 cells (Fig. 1F, Fig. S2I-J). In all, cFOOT-seq provides high-resolution insights into chromatin primary structure, encompassing chromatin accessibility, nucleosome positioning, and TF occupancy, even with minimal cell numbers, making it a powerful tool for studying chromatin dynamics and TF binding across diverse cell types and conditions.

### cFOOT-seq quantitatively measure TF occupancy

To comprehensively assess transcription factor occupancy and dynamics under various conditions through TF footprint analysis, we developed a computational framework named FootTrack (Footprint Analysis for Tracking TF Occupancy and Kinetics) (Fig. S3A), which conceptionally adapted from TOBIAS and Footprint tools^37, 40^. Using known TF binding information and motif information, FootTrack precisely maps TF occupancy and the chromatin landscape around TF motif centers at known binding sites. By integrating cFOOT-seq data and motif information from JASPAR^49^, FootTrack also facilitates de novo prediction of transcription factor binding sites genome-wide.

We quantified TF occupancy using the Footprint Occupancy Score (FOS), calculated as the difference between the average DNA conversion rate in the 50 bp flanking regions on either side of the motif and the motif center (Fig. 2A), similar to FOS calculation in DNase-seq^34^. Averaging FOS across all binding sites or specific regions yields Transcription Factor Occupancy Score (TFOS), which reflects general TF occupancy.

**Fig. 2.**
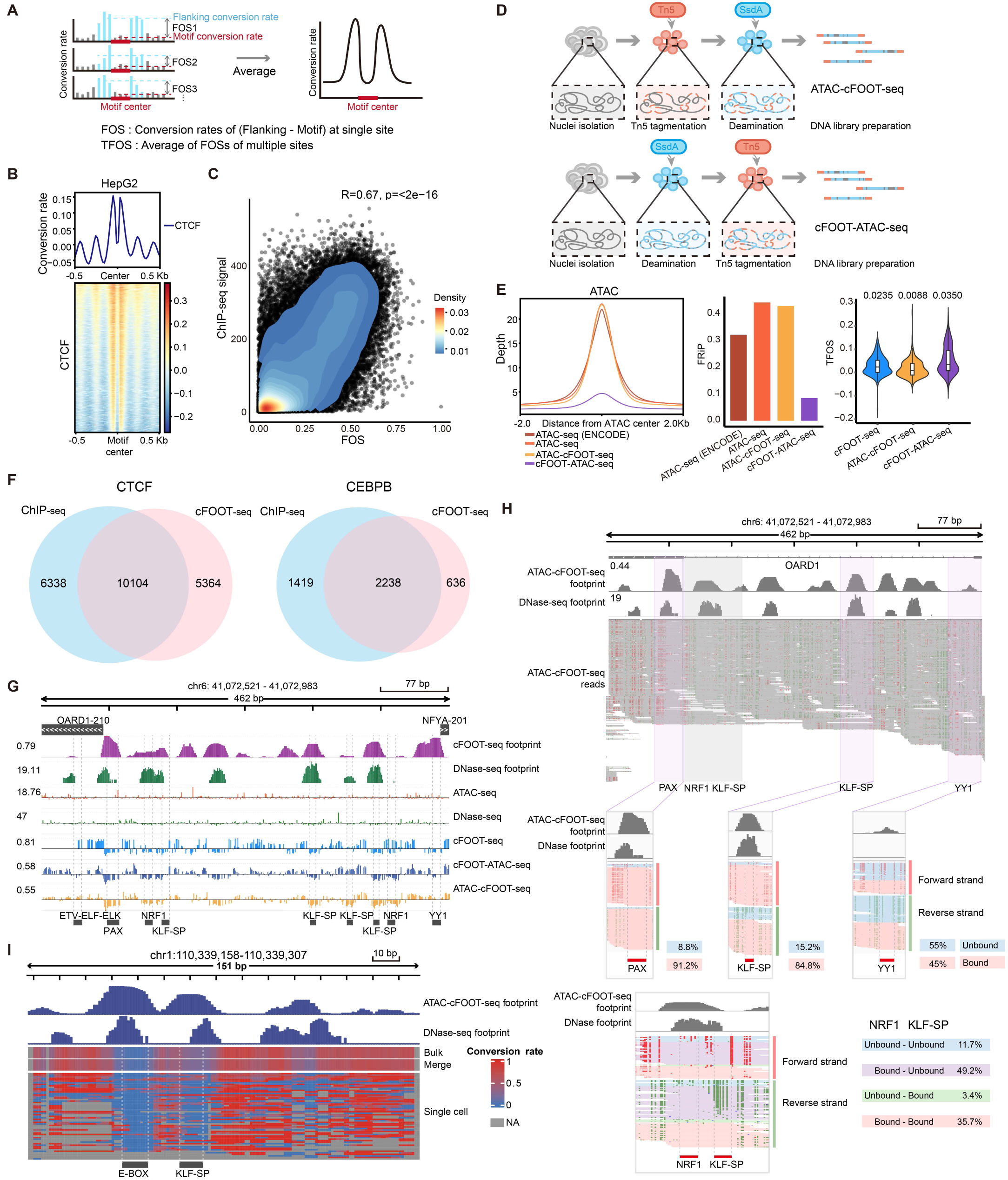
cFOOT-seq combined with ATAC-seq detects TF footprint at single-molecule and single-cell level. **A.** Schematic of calculations for Footprint Occupancy Score (FOS) and Transcription Factor Occupancy Score (TFOS). FOS is defined as the difference between the average conversion rate in the 50bp flanking regions on both side of the motif and the average conversion rate of the motif itself, representing the occupancy information at individual binding motif. TFOS is calculated as the average FOS across multiple binding sites of the same TF, reflecting the overall TF occupancy. **B.** Average profile of corrected DNA conversion rate around CTCF binding sites defined by ChIP-seq in HepG2 cells (30%). Heatmaps showing DNA conversion rate around individual CTCF binding sites aligned to motif centers and ranked by average conversion rate within ±0.5 kb of the motif center (bottom) around CTCF binding sites **C.** The scatter plot illustrates the Pearson correlation between cFOOT-seq and ChIP-seq signals at CTCF binding sites. cFOOT-seq signals are quantified using the Footprint Occupancy Score (FOS), including only sites with a FOS greater than 0. The ChIP-seq signal is calculated as the average value around each motif, extending ±50 bp from the motif’s center. The correlation coefficient is 0.67, with a highly significant p-value of less than 2e-16. The color gradient indicates the density of data points **D.** Workflow of cFOOT-ATAC-seq and ATAC-cFOOT-seq methods: cFOOT-ATAC-seq involves SsdA_tox_ deaminase incubation followed by Tn5-mediated tagmentation (top); ATAC-cFOOT-seq involves Tn5-mediated tagmentation followed by SsdA_tox_ deaminase incubation (bottom). **E.** Comparison of ATAC-seq, cFOOT-seq (30%), ATAC-cFOOT-seq (33%), and cFOOT-ATAC-seq (36%) methods: read depth around ±2 kb of the ATAC peak center based on 10 million reads, using two ATAC-seq datasets (one from ENCODE and one generated in-house) as benchmarks for evaluating open chromatin enrichment (Left), Fraction of Reads in Peaks (FRiP) (Middle) and distribution of TFOS (Right). **F.** Venn diagrams showing the overlap of CTCF and CEBPB footprints predicted by FootTrack-based de novo analysis of cFOOT-seq and CTCF binding motifs observed by ChIP-seq peaks in open chromatin regions of HepG2. **G.** IGV browser graphics showing profiles of cFOOT-seq, cFOOT-ATAC-seq, ATAC-cFOOT-seq, DNase-seq, and ATAC-seq at a representative site (chr6: 41,072,521–41,072,983) in HepG2 cells. The "cFOOT footprint" track shows footprint scores derived from cFOOT-seq data, while the "DNase footprint" track shows footprint probabilities from DNase-seq data. The five tracks below display the distribution of corrected cutting events for ATAC-seq and DNase-seq, along with the corrected conversion rate for cFOOT-seq, cFOOT-ATAC-seq, and ATAC-cFOOT-seq. The black labels at the bottom highlight motifs identified in ChIP-seq peaks, corresponding to TF binding sites **H.** IGV browser visualization showing the single-molecule profile of ATAC-cFOOT-seq at a representative site (chr6: 41,072,521–41,072,983) in HepG2 cells. The "ATAC-cFOOT-seq footprint" track represents footprint scores calculated from ATAC-cFOOT-seq data, while the "DNase footprint" track shows footprint probabilities predicted from DNase-seq data. The "ATAC-cFOOT-seq reads" track displays the individual mapped reads in this region. Both forward and reverse strands are shown for the three TF motifs (PAX, KLF-SP, and YY1), with reads categorized into bound and unbound states based on conversion rate at each motif. The bottom panel visualizes the distribution of reads for two adjacent motifs (NRF1 and KLF-SP), categorized into four groups based on the presence or absence of TF binding, providing insight into their single-molecule occupancy patterns. **I.** IGV browser visualization showing scATAC-cFOOT-seq profiles at a representative site (chr1:110,339,158–110,339,307) in K562 cells (38%). The “ATAC-cFOOT-seq footprint” track shows footprint scores calculated from bulk ATAC-cFOOT-seq data, while the “DNase footprint” track represents footprint probabilities from DNase-seq data. Below, conversion rate profiles for bulk (top), merged (middle), and single-cell (bottom) data are displayed, with the conversion rate color scale ranging from 0 (blue) to 1 (red). A 5-bp smoothing window is applied to the single-cell data, and sites with no data are marked in gray. The motifs E-BOX and KLF-SP, predicted in the footprint regions, are shown below the plot See also Figures S3, S4, S5, Table S1, S2

To optimize cFOOT-seq procedures and FootTrack analysis parameters, we selected HepG2 and K562 cell lines due to the extensive TF binding data available for these model cells. To assess potential biases in cFOOT-seq’s detection of TF footprints due to the sequence context preference of SsdA_tox_, we analyzed the footprint of the HepG2-specific factor HNF1B in K562 cells. We observed that HNF1B also exhibited clear footprints in K562, suggesting that SsdA_tox_’s sequence bias could distort quantitative assessments of TF occupancy (Fig. S3B). To address this, we treated naked genomic DNA with SsdA_tox_ and calculated the sequence context bias probability in human and mouse genomes (Fig. S3C). After correcting for this bias, the distortion in TF footprints was significantly reduced (Fig. S3B and S3C).

We further optimized the enzyme concentrations for cFOOT-seq. We selected 204 TFs with known localization and motif information in HepG2 cells (Table S1) for optimization testing and further analysis^50, 51^. HepG2 cells were treated with varying enzyme units, revealing that at conversion rate of 25% and 30%, both TFOS and the percentage of sites with positive FOS for each TF were similar (Fig. S3D). However, when conversion rate exceeded 40%, both TFOS and the percentage of sites with positive FOS decreased. Therefore, we determined that a conversion rate range of 25-40% is optimal for accurately measuring most TFs. Next, we assessed the required sequencing depth for accurate FOS measurement. We found that a minimum of approximately 15 million reads (∼1.4x depth) per whole-genome sequencing sample is necessary for TFOS assessment, with approximately 200 million paired reads (∼6.5x depth) providing good stability and 400-800 million reads offering even higher reliability (Fig. S3E).

With these optimizations, FootTrack accurately depicted nucleosome organization around CTCF binding sites and measured CTCF occupancy in HepG2 cells using cFOOT-seq data (Fig. 2B). By calculating the FOS for CTCF binding motifs, we observed strong correlations between FOS and ChIP-seq signals (Fig. 2C), suggesting that FOS can serve as a reliable quantitative measure of transcription factor occupancy.

### cFOOT-seq combined with ATAC-seq provides high-resolution and sensitive detection of TF footprint

Compared to DNase-seq and ATAC-seq, cFOOT-seq converts chromatin structure information into DNA sequence data without disrupting chromatin structure, allowing combination with other chromatin probing technologies. We explored combining cFOOT-seq with ATAC-seq to enrich accessible chromatin and measure TF occupancy at a lower sequencing cost. We tested two combined strategies based on the order of treatment of SsdA_tox_ and Tn5: (1) ATAC-cFOOT-seq (Tn5 followed by SsdA_tox_) and (2) cFOOT-ATAC-seq (SsdA_tox_ followed by Tn5) (Fig. 2D).

ATAC-cFOOT-seq efficiently enriches open regions similar to ATAC-seq and detects footprints for most TFs, with TFOS generally comparable to those identified by cFOOT-seq (Fig. 2E, Fig. S4A-B). However, for certain TFs such as FOXA1, their TFOS were reduced (Fig. S4B), likely due to TF binding loss during the ATAC-cFOOT-seq process. cFOOT-ATAC-seq also demonstrates obvious enrichment of open regions compared to cFOOT-seq (Fig. S4C), though weaker than cFOOT-ATAC-seq (Fig. 2E). Importantly, cFOOT-ATAC-seq detected higher TFOS than cFOOT-seq, suggesting its greater stability and sensitivity in detecting TF binding (Fig. 2E, Fig. S4A-B) We noticed that Fraction of Reads in Peaks (FRiP) of cFOOT-ATAC-seq is negatively correlated with the conversion efficiency (Fig. S4D). When conversion efficiency is between 20-30%, cFOOT-ATAC-seq achieves higher enrichment of open regions while maintaining sensitive TF binding detection (Fig. S4D-E). In summary, ATAC-cFOOT-seq and cFOOT-ATAC-seq each offer distinct advantages in detecting TF footprints and enriching open chromatin regions, making them powerful and complementary techniques for TF footprint analysis DNase-seq has long been a benchmark for TF footprint detection^32, 34, 35, 40^, and ATAC-seq, with optimized bioinformatics is becoming increasingly popular^36, 37, 39^. By analyzing metrics including FWHM and Integral Width, we compared cFOOT-seq and its combined methods to DNase-seq and ATAC-seq for footprint resolution of 204 TFs in HepG2. While DNase-seq outperforms ATAC-seq in some instances, our results show that cFOOT-seq and its combined methods, particularly cFOOT-ATAC-seq, offer higher resolution overall (Fig. S4F).

Using TFs such as ATF4, CEBPD, HNF1A, and FOXA1, we further evaluated the resolution and sensitivity of these methods at different sequencing depths. At 13.2 M and1.65 M reads, ATAC-seq struggles to detect footprints, underscoring its dependence on sequencing depth (Fig. S4G). In contrast, cFOOT-seq and DNase-seq perform well, with cFOOT-seq detecting sharper footprints for ATF4, CEBPD, and HNF1A (Fig. S4G). Notably, while DNase-seq struggles with FOXA1, cFOOT-seq reliably detects its footprints. At extremely lower sequencing depths (0.33M reads), both DNase-seq and cFOOT-seq show decreased stability in footprint detection (Fig. S4G). However, the combined methods (ATAC-cFOOT-seq and cFOOT-ATAC-seq) maintain robust detection for all four TFs (Fig. S4G). This demonstrates the advantage of combining cFOOT-seq with ATAC-seq, which not only enriches open chromatin regions but also utilizes deaminase-mediated DNA conversion for more reliable and sensitive footprint detection, even at reduced sequencing depths.

To de novo identify genome-wide TF footprints from DNA conversion data of cFOOT-seq and investigate TF binding dynamics, FootTrack was used to predict the footprints in open chromatin of the whole genome. FootTrack offers two background correction modes: global background and local background. Following the background correction, FootTrack proposed two different strategies for predicting footprints: S1, calculates footprint scores for each motif with bias corrected data, and applies statistical methods to identify motifs with higher scores as potential TF binding sites, and S2, initially detects footprint regions by binomial statistical test and then scans for motifs within the identified footprint regions (Fig. S3A). Based on the performance evaluation, we found that local mode performs better than global mode for both S1 and S2, and S1 performed better for cFOOT-seq, while S2 performs better for ATAC-cFOOT-seq and cFOOT-ATAC-seq (Fig. S4H).

To validate the accuracy of footprint prediction, we examined representative TFs with well-characterized ChIP-seq binding profiles. The footprints of CTCF and CEBPB identified by FootTrack in open chromatin regions showed strong overlap with motifs located within ChIP-seq peaks in HepG2 (Fig. 2F). We further applied FootTrack to cFOOT-seq and cFOOT-ATAC-seq data from HepG2, K562, and R1 cells to quantify footprint density in open chromatin regions (OCRs). Footprint density was similar across cell types (∼4.2 footprints per 200 bp; Fig. S4I–L). Notably, enhancer-associated OCRs consistently showed higher footprint density than promoter-associated OCRs, suggesting more extensive TF binding at enhancers^40^. To assess how many footprints could be linked to known TFs, we compared predicted footprints in open regions with known motifs. About 60% matched known motifs, while ∼40% lacked recognizable matches (Fig. S4L–N). Although footprint scores were similar, footprints without known motifs were generally broader, potentially reflecting binding by larger chromatin-associated proteins, TF complexes, or cooperative TF interactions. Notably, an increasing number of TF pairs have been found to form composite motifs distinct from their individual canonical motifs^52, 53^. Since our analysis primarily relied on the JASPAR database, incorporating additional motif sources and accounting for composite motif architectures in future work may further improve the annotation of currently unassigned footprints.

### cFOOT-seq combined with ATAC-seq enables detection of TF footprint at single-molecular and single-cell level

To study transcription factor function, accurately detecting TF occupancy at individual sites is crucial for footprint-based methods. Using cFOOT-seq and its combined methods, we analyzed two regions (chr6: 41,072,521–41,072,983 and chr1:110,338,780-110,339,310) and found that cFOOT-seq and its combined methods predict footprints that overlap with those defined by DNase-seq and motifs in ChIP-seq peaks, demonstrating their reliability in detecting TF footprints at these sites in a de novo manner (Fig. 2G, Fig. S5A).

Combining cFOOT-seq with ATAC-seq enriches open chromatin regions, enabling higher sequencing depth and more precise quantification of TF occupancy at both single-molecule and single-cell levels. At the single-molecule level, TF occupancy is quantified by analyzing conversion rate at footprint sites, with each read covering the corresponding motif. For example, at region 1, variations in TF occupancy across different motifs are observed: PAX motif (ATAC-cFOOT-seq, 91.2%; cFOOT-ATAC-seq, 98%), KLF-SP motif (ATAC-cFOOT-seq, 84.8%; cFOOT-ATAC-seq, 76.9%), and YY1 motif (ATAC-cFOOT-seq, 45%; cFOOT-ATAC-seq, 69.4%) (Fig. 2H, S5B).

Similar analyses for the two E-box motifs and ZBTB33 at region 2 further confirm this capability (Fig. S5C-D). These findings underscore cFOOT-seq’s ability to capture detailed TF occupancy insights through conversion patterns in footprint reads. Additionally, single-molecule conversion profiles allow the analysis of TF occupancy at nearby sites, such as NRF1 and KLF-SP motifs in region 1 (Fig. 2H). Notably, 49.2% of cases show NRF1 bound and KLF-SP unbound, 35.7% show both NRF1 and KLF-SP bound together, and only 3.4% show KLF-SP bound without NRF1, suggesting a potential dependence of KLF-SP binding on NRF1 at this site. This ability to detect TF co-occupancy within the same read forms the basis for studying synergistic binding events between TFs.

By integrating plate-based single-cell library preparation technology, we enable TF footprint detection at the single-cell level through scATAC-cFOOT-seq. After Tn5 tagmentation and SsdA_tox_ deamination, cells are sorted into individual wells of a 96-well plate. Following cell lysis, DNA is barcoded by preamplification using well-specific primers, and the samples are pooled for further library amplification (Fig. S5E). A total of 71 barcodes (Table S2) were used to label 71 K562 cells, and 100 million reads were sequenced (∼1.3 million reads per cell). On average, we detect over 25,000 peaks per cell, with FRiP greater than 0.4 (Fig. S5F). TF footprint signals at the binding sites of CTCF and NRF1 were observed in individual cells (Fig. S5G). Notably, TF footprints can even be detected at single genomic loci within individual cells (Fig. 2I), demonstrating the ability to analyze TF occupancy in heterogeneous cell populations. Together, cFOOT-seq combined with ATAC-seq provides a high-resolution, powerful tool for detecting TF footprints at both the single-molecule and single-cell levels, offering valuable insights into gene regulatory networks, TF interactions, and chromatin dynamics.

### cFOOT-seq quantitatively assess the impact of chromatin accessibility, histone modifications, and cofactors on TF occupancy

TF binding is influenced by chromatin accessibility and histone modifications, but large-scale systematic analysis is limited. Now cFOOT-seq provides us a chance to quantitatively analyze factors associated with TF binding.

To investigate the impact of chromatin accessibility on TF occupancy, we analyzed 204 TFs in HepG2 and 127 TFs in K562 with known binding sites and motifs (Table S1). We categorized their binding sites into four groups: closed chromatin regions (CCR), and low, moderate, and high open chromatin regions (OCR). Our analysis revealed a general trend where higher chromatin accessibility correlated with increased TFOS (Fig. S6A and S6B). While many TFs displayed higher TFOS in highly accessible regions, some TFs with lower overall TFOS showed no significant change across different levels of chromatin accessibility (Fig. S6A).

We further selected 89 overlapping TFs from both cell lines and clustered them based on their TFOS in CCR and OCR (Fig. 3A). The differences in TFOS between closed and open regions were generally consistent between HepG2 and K562 (Fig. S6C), indicating that chromatin accessibility similarly influences TF occupancy in both cell types. Among the 89 TFs analyzed, CTCF, ZNF24, ZBTB33, TCF7, USF2, CEBPB, and FOX family proteins exhibited high TFOS (>0.05) in CCR in both cell types (Fig. 3B). This indicates these proteins have a stronger ability to bind closed chromatin. Consistently, FOX proteins have been proposed as pioneer factors capable of accessing closed chromatin. Additionally, CTCF, bHLH E-box binding factors (USF2, TFE3, MLX, BHLHE40), bZIP factors (CREB1, CREM, ATF3/7), and RFX family proteins (RFX1/5) demonstrated high TFOS in OCR in both cell lines, with TFOS in open regions being significantly higher than in closed regions (Fig. 3A, 3C and S6A). This suggests their preferential binding to accessible chromatin.

**Figure 3:**
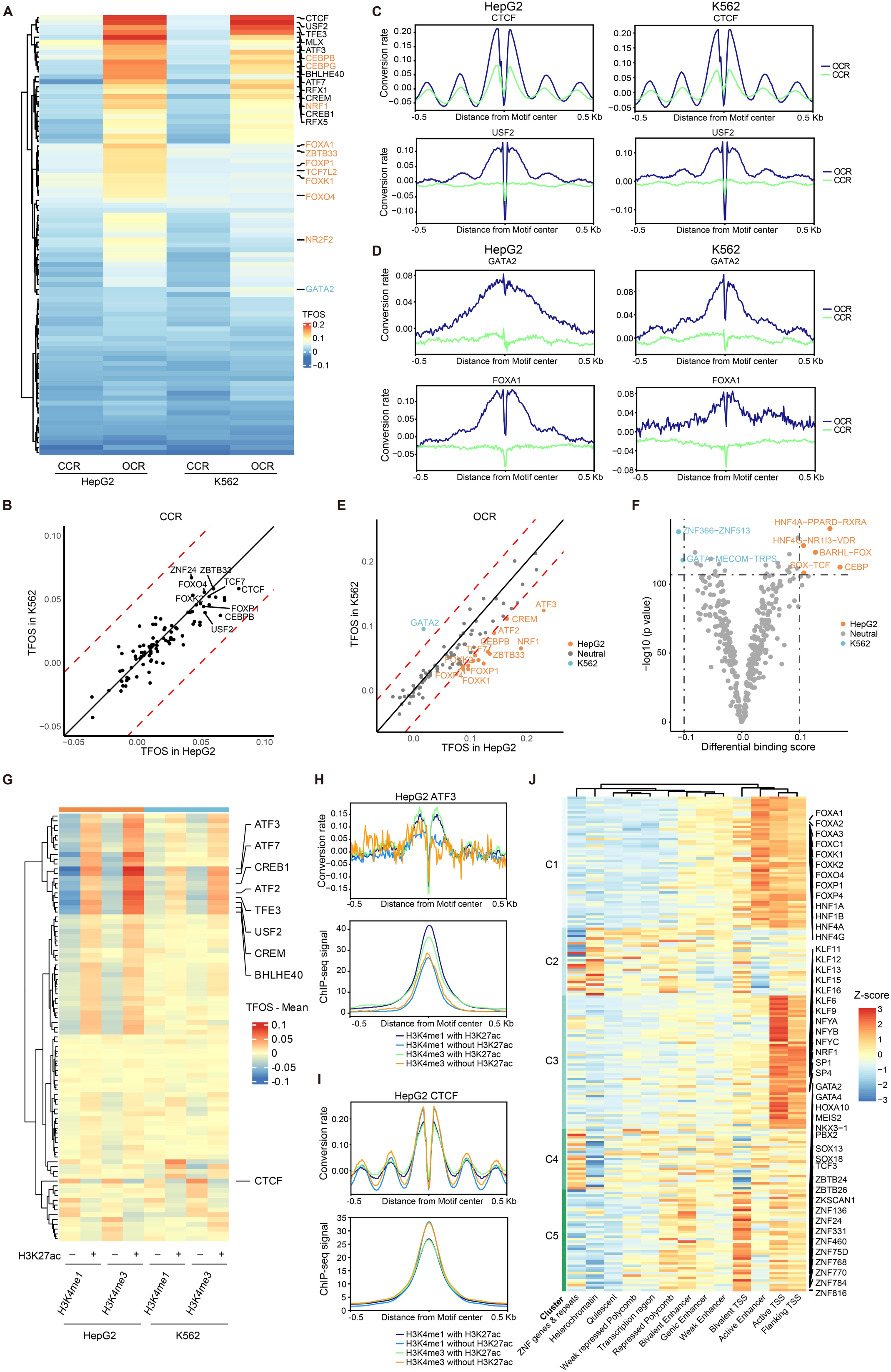
cFOOT-seq quantitatively assess the impact of chromatin accessibility, histone modifications, and cofactors on TF occupancy. **A.** Heatmap of TFOS at closed chromatin regions (CCR) and open chromatin regions (OCR) of 89 overlapping TFs with known binding sites and motifs in HepG2 (30%) and K562 (30%) cells. CCR and OCR are defined by ATAC-seq in each cell type. TFs highlighted in orange indicate those with higher TFOS in OCR of HepG2, while TFs highlighted in blue indicate those with higher TFOS in OCR of K562. The color scale represents TFOS depth, with higher values in red and lower values in blue. **B.** Scatter plot comparing the TFOS in CCR between HepG2 and K562 cells. Each point represents an individual transcription factor. TFs with TFOS (in CCR) >0.05 in both cells are highlighted. The red dashed lines denote a threshold where the difference in TFOS between the two cell types exceeds 0.04. **C-D.** Average profiles of corrected DNA conversion rate around the binding sites of CTCF and USF2 (C), GATA2 and FOXA1 (D) in HepG2 and K562 cells, highlighting their differences of TFOS in CCR and OCR, and TFOS difference between cell lines. **E.** Scatter plot comparing the TFOS of TFs at known binding sites in OCR between HepG2 and K562 cells. The red dashed lines denote a threshold where the difference in TFOS between the two cell types exceeds 0.04. TFs highlighted in orange represent those with TFOS values in HepG2 that are greater than in K562 by more than 0.04, while TFs highlighted in blue represent those with TFOS values in K562 that are greater than in HepG2 by more than 0.04. **F.** Volcano plot showing the TF clusters with differential binding scores between HepG2 and K562 cells, based on de novo binding sites predicted by FootTrack. TFs with significantly higher binding score in HepG2 are colored in orange, and those with higher binding score in K562 are colored in blue. **G.** Heatmap showing TFOS in regions with different histone modification status (H3K4me1^+^ H3K27ac^-^, H3K4me1^+^ H3K27ac^+^, H3K4me3^+^ H3K27ac^-^, and H3K4me3^+^ H3K27ac^+^) of 89 overlapping TFs in HepG2 and K562 cells. **H-I.** Average profiles showing corrected DNA conversion rate (top) and ChIP-seq signal (bottom) around ATF3 (H) and CTCF (I) binding sites in HepG2 cells. Binding sites were defined by ChIP-seq of each TF. Both cFOOT-seq and ChIP-seq signals are shown separately for regions marked by different combinations of histone modifications (H3K4me1^+^ H3K27ac^-^, H3K4me1^+^ H3K27ac^+^, H3K4me3^+^ H3K27ac^-^, and H3K4me3^+^ H3K27ac^+^) of HepG2 cells. **J.** Heatmap showing TFOS Z-score in regions defined by chromHMM annotations for 204 TFs in HepG2 cells. The TFs were clustered into five groups, with the representative TF highlighted. See also Figures S6, Table S1, S3

Comparing TFOS between HepG2 and K562 cells revealed potential cell-specific transcription factors. While TFOS in CCR was generally similar between the two cell types, notable differences were observed in OCR (Fig. 3B, 3D-E). HepG2 cells exhibited higher TFOS for factors such as NRF1, NR2F2, TCF7L2, ZBTB33, bZIP family members (ATF3, CEBPB/CEBPG, CREM), and FOX family proteins, whereas K562 cells showed higher TFOS for GATA2 (Fig. 3D-E). To further identify cell-specific factors, we utilized FootTrack for de novo footprint analysis. This analysis predicted higher scores for clusters such as CEBP, BARHL-FOX, HNF4A-PPARD-RXRA, SOX-TCF, in HepG2, and for the GATA cluster in K562 (Fig. 3F). These predictions were consistent with the cFOOT-seq signal at known TF binding sites derived from ChIP-seq (Fig. 3E and S6D), demonstrating the predictive capability of FootTrack.

To investigate how co-binding influences TF occupancy, we used TF-COMB^54^ to identify co-occurring TF pairs in open chromatin regions based on ChIP-seq data for 204 TFs in HepG2, 127 in K562, and 27 in R1. Top-ranked pairs such as ZBED4–EGR1 (HepG2, Fig. S6E) and SP1–MAZ (K562, Fig. S6F) showed enhanced TFOS in shared regions compared to individual binding sites, and both TF pairs showed higher chromatin accessibility in co-bound regions, suggesting that TF co-occupancy may promote chromatin accessibility and enhance TF binding. Further Motif spacing analysis revealed ZBED4–EGR1 displayed a broader spacing distribution, suggesting less spatial constraint and flexible chromatin co-binding (Fig. S6E), while SP1–MAZ showed a strong bias toward near-zero spacing (Fig. S6F), likely due to motif similarity and overlapping binding, which may cause steric hindrance. Thap11 and ZNF143 were identified as a prominent co-binding pair in R1 cells. Their motif centers were enriched at an 8 bp spacing, consistent with previous reports of composite motif formation^55^ (Fig. S6G). Notably, co-occupancy markedly increased ZNF143’s TFOS, while Thap11 was only modestly affected (Fig. S6G), indicating that ZNF143 may depend on Thap11 for stable chromatin association. Together, these findings suggest that cooperative binding and increased chromatin accessibility are key contributors to enhanced TFOS in shared regions, while steric hindrance may also play a role in specific TF combinations.

To investigate the effect of histone modifications on TF occupancy, we analyzed the TFOS of 89 TFs in HepG2 and K562 across four chromatin states: H3K4me1^+^ H3K27ac^-^, H3K4me1^+^ H3K27ac^+^, H3K4me3^+^ H3K27ac^-^, and H3K4me3^+^ H3K27ac^+^ (Fig. 3G). Generally, TFOS was higher in H3K27ac+ regions compared to H3K27ac^-^ regions. Specifically, bHLH E-box binding factors (USF2, TFE3, BHLHE40) and bZIP factors (CREB1, CREM, ATF2/3/7) exhibited significantly higher TFOS in H3K27ac^+^ regions than in H3K27ac^-^ regions, regardless of H3K4me1 or H3K4me3 presence, consistent with the trends observed in ChIP-seq data (Fig. 3G-H, S6H). Moreover, the TFOS of CTCF was notably lower in H3K27ac^+^ regions compared to H3K27ac^-^ regions (Fig. 3I, S6I). This trend was also evident in ChIP-seq data, suggesting a negative correlation between CTCF binding and the presence of H3K27ac or its associated proteins.

To comprehensively analyze the impact of chromatin context on TF occupancy, we clustered 204 TFs in HepG2 cells into five groups based on their TFOS in regions defined by chromHMM annotations^56^ (Fig. 3J, and Table S3). We found that Cluster 1, enriched with liver-specific factors and FOX family proteins, had the highest TFOS in active enhancer regions, consistent with their role as tissue-specific pioneer factors that open chromatin at enhancers to regulate liver-specific genes. Cluster 3 was enriched with promoter-binding factors, which had the highest TFOS at transcription start sites (TSS) and TSS-flanking regions. Cluster 5 included TFs involved in embryonic development and cell fate determination, as well as ZNF proteins, which had the highest TFOS in bivalent TSS regions. This indicates that TFs tend to have higher binding affinity in their preferred regions, where they play key functional roles, suggesting significant binding specificity and functional specialization of TFs. The specialization of TFs to distinct genomic contexts underscores the intricate nature of TF-chromatin interactions and their critical role in shaping the regulatory landscape of the genome.

### cFOOT-seq could capture the dynamics of TF in early stage of OSKM reprogramming

Unraveling core transcription factor regulatory networks is crucial for understanding cell identity during differentiation and reprogramming^33^. Using cFOOT-seq, we analyzed TF binding differences between MEF and R1 cells. By inputting the open chromatin regions of MEF and R1 into FootTrack, we identified TFs enriched in each cell type (Fig. S7A). Further analysis with TF-COMB revealed that AP-family TFs form a closely co-occurrence network in MEF, while stemness-related factors such as SOX family proteins and SALL4 form a network in R1 (Fig. S7B). These results suggest that cFOOT-seq effectively captures cell type-specific TFs and provides insights into TF co-occurrence network.

The reprogramming of MEF to iPSC is coupled with dramatic changes of chromatin accessibility and TFs binding^57, 58^. To assess if cFOOT-seq could capture TF dynamics during reprogramming, we used an inducible OSKM system to reprogram MEF to iPS cells. Doxycycline (Dox) was added for 24, 48, and 96 h to induce OSKM expression, and TF footprints were detected using cFOOT-seq (Fig. 4A). TFOS of OCT4 and KLF4 increased rapidly after 24 h of induction (Fig. 4B). Conversion rate at the flanking sites of the OCT4 binding motif increased after Dox addition, while the rate at the flanking sites of the KLF4 motif remained high and unchanged (Fig. 4B). This suggests that OCT4 acts as a pioneer factor opening chromatin at closed sites, whereas KLF4 binds to pre-accessible chromatin in MEF post-induction.

**Fig. 4.**
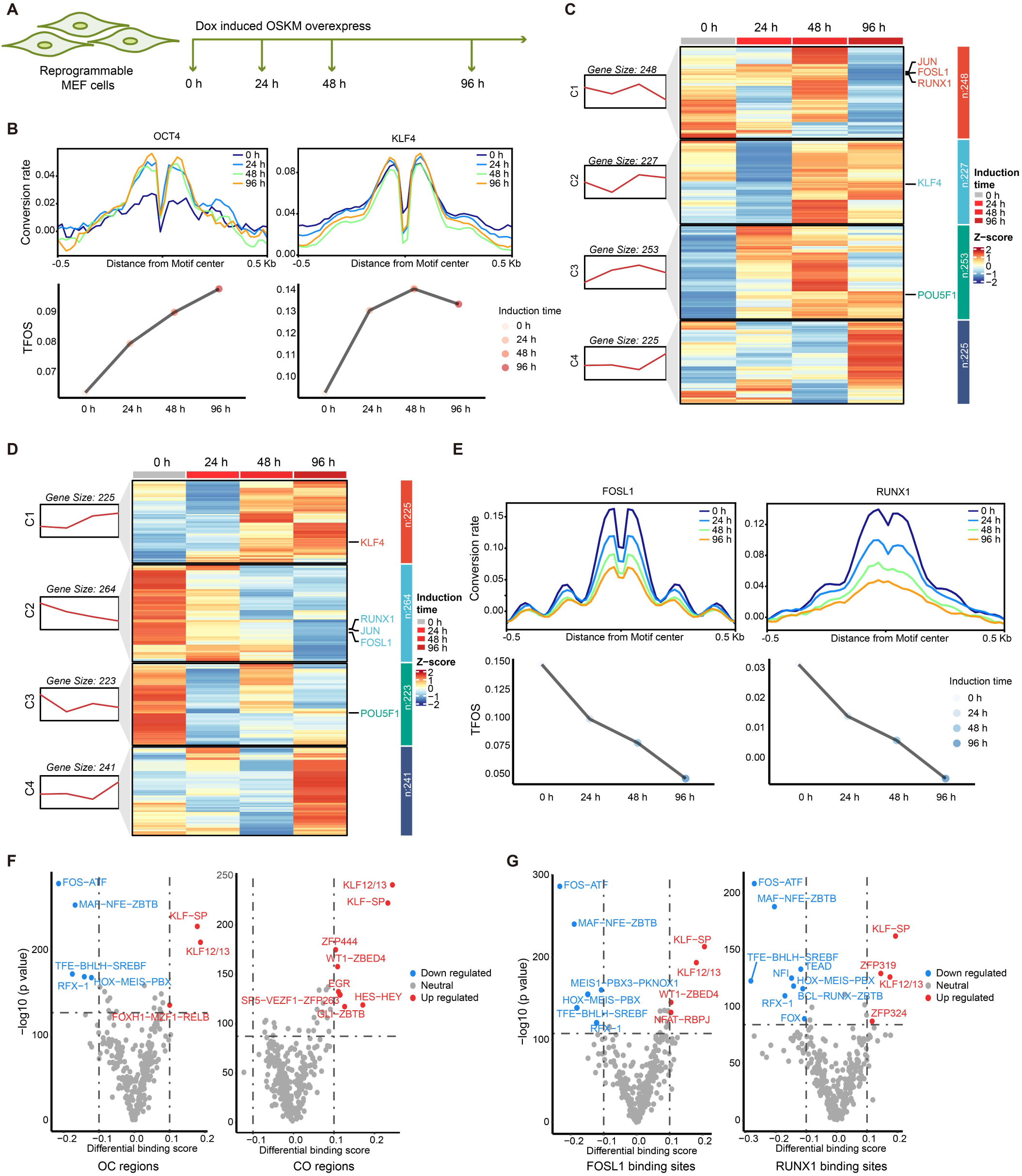
cFOOT-seq could capture the dynamics of TFs in early stage of OSKM reprogramming. **A.** Schematic of experimental treatments on reprogrammable MEF cells with doxycycline-induced OSKM overexpression. **B.** Average profiles and box plots showing changes in OCT4 (left) and KLF4 (right) footprints over induction times (0 h-34%, 24 h-34%, 48 h-35%, 96 h-37%) in OSKM-MEF cells. Average profiles display corrected DNA conversion rate aligned at OCT4 and KLF4 binding sites in R1. Line plots show TFOS changes representing dynamics of TF occupancy. **C.** Heatmaps of FootTrack-predicted TF footprints in R1-specific open regions at different induction times (0h, 24h, 48h, 96h), clustered into four groups by footprint changes. **D.** Heatmaps of FootTrack-predicted TF footprints in MEF-specific open regions at different induction times (0 h, 24 h, 48 h, 96 h), clustered into four groups by footprint changes. **E.** Average profiles and box plots showing changes in FOSL1 and RUNX1 footprints over time in OSKM-MEF cells. Average profiles display corrected DNA conversion rate aligned at FOSL1 and RUNX1 known binding sites defined by ChIP-seq in R1. Line plots show TFOS changes representing dynamics of TF occupancy. **F.** Volcano plots showing the FootTrack-predicted TF clusters with differential binding scores between 0 h and 48 h in OSKM-MEF cells for OC regions (open in MEF, closed in R1) and CO regions (closed in MEF, open in R1). TFs with upregulated footprint scores at 48h are highlighted in red; TFs with downregulated scores are highlighted in blue. **G.** Volcano plots showing the FootTrack-predicted TF clusters with differential binding scores between 0 h and 48 h in OSKM-MEF cells for FOSL1 and RUNX1 binding regions in MEF cells. TFs with upregulated footprint scores at 48h are highlighted in red; TFs with downregulated scores are highlighted in blue. See also Figures S7

To systematically analyze TF changes, we used FootTrack to predict TF occupancy dynamics in MEF-specific and R1-specific open regions during early reprogramming, clustering TFs based on TFOS variation trends (Fig. 4C-D). TFOS for JUN, FOSL1, and RUNX1 decreased significantly in MEF-specific regions (Fig. 4C), while KLF4 and POU5F1 increased in both MEF and R1-specific open regions (Fig. 4D). The TFOS decrease of FOSL1 and RUNX1 was confirmed by the cFOOT-seq signal at known TF binding sites defined by ChIP-seq (Fig. 4E), supporting that the binding of these proteins is decreased during the reprogramming of MEF to R1. TF-comb analysis has revealed that AP family TF interactions are more specific to MEF (Fig. S7B), the reduced AP family binding is required for MEF reprogramming towards iPS cells, consistent with the idea that AP family proteins act as reprogramming barriers^58, 59^.

Based on differential chromatin accessibility between MEF and R1, we defined OC regions as those open in MEF but closed in R1, and CO regions as those closed in MEF but open in R1. We further analyzed the potential driving forces behind changes in chromatin accessibility during reprogramming and found that KLF family proteins increased in both OC and CO regions (Fig. 4F). These results suggest that increased binding of KLF proteins may be correlated with changes of chromatin accessibility during reprogramming. Additionally, further analysis of the binding sites of FOSL1 and RUNX1 in OC regions indicated that KLF proteins might be associated with the suppression of FOSL1 and RUNX1 (Fig. 4G) ^59^, highlighting their role in gene regulation during the reprogramming process.

### cFOOT-seq depicts dynamics of nucleosome organization and TF occupancy in response to inhibition of SWI/SNF

Chromatin remodeling complexes regulate gene expression by modulating the organization of nucleosome through ATP-dependent translocase activity^60-63^. SWI/SNF remodeler could open chromatin by nucleosome sliding and eviction, facilitate the binding of TFs, and its mutations or dysfunction are linked to various cancers and developmental disorders^64-67^. Although there have been efforts to explore the effect of SWI/SNF complex on chromatin accessibility and the binding of TFs^68-73^, a comprehensive understanding of TF dependence on SWI/SNF remains limited due to the lack of high-throughput methods for accurately and quantitatively measuring TF occupancy on chromatin. To address this, we used cFOOT-seq to study chromatin accessibility around TFs and dynamics of TF occupancy following SWI/SNF inhibition with compound BRM014 in mESC cells and HepG2 cells^74^.

In mESC treated with BRM014 for 1 and 24 h, we observed a general decrease in conversion rate around ATAC-seq peaks, indicating reduced chromatin accessibility (Fig. S8A-B). Consistent with previous reports, TFs such as OCT4 and REST showed significant decreases in DNA conversion rate around the binding sites of OCT4 and REST, along with a reduction in TFOS for these factors, while CTCF binding sites remained unchanged (Fig. S8C)^69^. FootTrack predicts that TF clusters, including GLI-ZIC, TFE-BHLH-SREBF, bHLH-PAS, ESRR-NR, KLF, MYC proteins, had decreased footprint scores (Fig. S8D). Published ChIP-seq data confirmed that the footprints of MYCN and KLF4 decreased after BRM014 treatment (Fig. S8E). However, the chromatin accessibility pattern around the binding sites of MYCN and KLF4 largely remained unchanged (Fig. S8E).

To comprehensively analyze the effect of SWI/SNF inhibition on nucleosome organization and TF occupancy, we selected HepG2 cells as our model system due to the extensive TF binding data available from the ENCODE project. HepG2 cells were treated with the SWI/SNF inhibitor BRM014 for 1, 6, and 24 h, followed by recovery periods of 6 and 24 h after the 24 h treatment (Fig. 5A). Similar to the observations in R1 cells, the conversion rate around ATAC-seq peaks in HepG2 decreased rapidly in response to BRM014 treatment and recovered upon inhibitor removal (Fig. S8F). ATAC-seq peaks overlapping with the SWI/SNF complex showed a more dramatic decrease compared to those without SWI/SNF, consistent with the inhibitory effect of BRM014 on the SWI/SNF complex (Fig. 5B).

**Figure 5:**
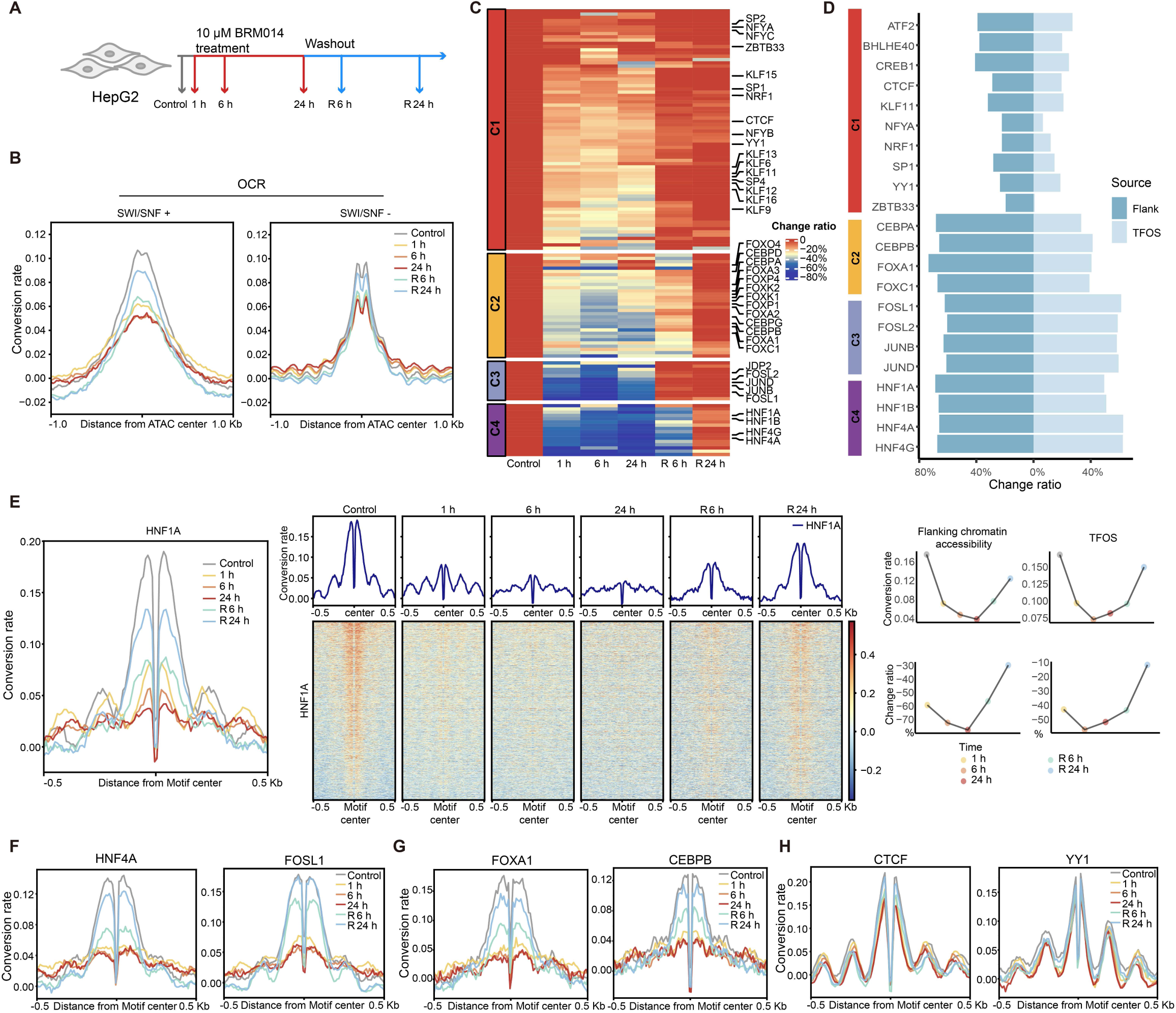
cFOOT-seq depicts dynamics of nucleosome organization and TF occupancy in response to inhibition of SWI/SNF. **A.** Schematic representation of the experimental design depicting the treatment of HepG2 cells with 10 μM BRM014 for 1, 6, and 24 h, as well as 6 h and 24 h recovery after 24 h BRM014 treatment. **B.** Average profiles of DNA conversion rate around OCR with or without SWI/SNF binding sites defined by ChIP-seq, illustrating changes in chromatin accessibility following different durations of BRM014 treatment and recovery (Control 0 h-32%, 1 h-31%, 6 h-32%, 24 h-29%, R6 h-33%, R24 h-33%). **C.** Heatmap showing TFOS change ratio in SWI/SNF+ regions across different BRM014 treatment conditions for 134 filtered TFs in HepG2 cells. The TFs are clustered into four groups based on change ratio of TFOS across treatments. Grids color represents the dynamic change ratio compared with controls **D.** Butterfly graph shows flanking chromatin accessibility change ratio (left) and TFOS change ratio (right) of representative TFs in different clusters, which represent changes in flanking accessibility and TF occupancy. **E.** Average profiles and heatmap of overall DNA conversion rate around HNF1A binding sites in SWI/SNF+ regions (left), illustrating changes in chromatin accessibility, nucleosome position and TF occupancy after BRM014 treatment and recovery. Plot graphs of flanking chromatin accessibility, TFOS and their change ratio are shown (Right). Flanking chromatin accessibility around the TF binding motifs are calculated as the average DNA conversion rate in the 50 bp flanking regions on both sides of the motif as flanking chromatin accessibility **F-H.** Average profiles of DNA conversion rate around TF binding sites of HNF4A and FOSL1 (F), FOXA1 and CEBPB (G), CTCF and YY1 (H) under control, 1 h, 6 h, and 24 h BRM014 treatments, as well as 6 h and 24 h recovery after 24 h BRM014 treatment. The profiles show changes in chromatin accessibility and TF footprints across different treatment conditions. See also Figures S8, Table S4

To accurately assess changes in TF occupancy, we calculated the TFOS at SWI/SNF+ sites for individual TFs and determined the TFOS change ratio post-treatment. To ensure reliability, we filtered out TFs with unrobust TFOS as mentioned in methods, resulting in a selection of 134 qualified TFs. We then profiled their dynamic responses to BRM014 treatment by clustering TFs based on their TFOS change ratios (Fig. 5C, Table S4). To measure chromatin accessibility flanking the TF binding sites, we calculated the average DNA conversion rate in the 50 bp flanking regions on both sides of the motif as flanking chromatin accessibility. Based on these clusters, we further calculated the change ratio of flanking chromatin accessibility post-treatment (Fig. S8G). Representative TFs in clusters 3 and 4 showed a significant decrease in both flanking chromatin accessibility and TFOS following treatment, with TFs in cluster 3 recovering more quickly than those in cluster 4(Fig. 5C-D, S8G). TFs in cluster 2 exhibited a marked decrease in flanking chromatin accessibility but only a mild reduction in TFOS. Conversely, TFs in cluster 1 demonstrated minimal decreases in both flanking chromatin accessibility and TFOS (Fig. 5C-D, S8G).

We further analyzed the conversion rate kinetics around TF binding sites of representative TFs to assess dynamics of flanking chromatin accessibility and nucleosome organization. For cluster 4, HNF1A at SWI/SNF+ sites showed a dramatic decrease in TFOS and conversion rate of flanking chromatin around its binding sites after 1 h of BRM014 treatment, with nucleosome positions moving closer to the binding sites (Fig. 5E). This suggests a quick loss of HNF1A binding, chromatin accessibility and nucleosome repositioning. After 6 and 24 h of treatment, the nucleosome phasing pattern further diminished, likely due to the loss of the anchoring effect caused by HNF1A binding. Similar to HNF1A, decrease of flaking chromatin accessibility and TF binding were observed for HNF1B/4A/4G in cluster 4, and AP-1 family proteins (FOSL1/2, JUNB/D) in cluster 3 which are highly sensitive to BRM014, suggesting the chromatin accessibility around the binding motifs and TF occupancy of these proteins are largely dependent on SWI/SNF (Fig. 5E-F, S8H).

For cluster 2, the conversion rate around the binding sites of FOX family are also decreased dramatically, suggesting the chromatin accessibility around their binding sites are dependent on SWI/SNF. However, compared to HNF proteins and AP-1 family proteins, the TFOS change ratios of FOX family proteins are lower (Fig. 5D, 5G), indicating that their binding on chromatin is less dependent on the opening of the chromatin by SWI/SNF, which is also consistent with the pioneer activity of FOX proteins to bind closed chromatin.

For cluster 1, CTCF showed only mild decreases in TFOS and chromatin accessibility around their binding sites (Fig. 5H). Consistent with this observation, NURF-specific subunit BPTF and ISWI core translocase SNF2H has been reported to maintain the organization of nucleosome and local accessibility around CTCF sites ^75-77^. Additionally, NRF1, NFY complex, YY1, and ZBTB33 were resistant to BRM014 treatment (Fig. 5H, S8I). It’s highly possible that like CTCF, these proteins may also dependent on other remodeling factors to open chromatin, which need further investigation^75, 78^.

### Definition of TF dependence on SWI/SNF reveals the spatial organization rule of TFs

Previous reports have shown that chromatin accessibility at promoters and enhancers may respond differently to the deletion or inhibition of SWI/SNF ^71, 73, 79^. We also found that compared to promoters, the overall conversion rate of enhancers decreased more obviously after BRM014 treatment in HepG2, suggesting that the chromatin accessibility of enhancers are more dependent on SWI/SNF complex (Fig. S9A). To evaluate the differential effect of enhancers and promoters on TF dependence of SWI/SNF, we separated the TF binding sites by promoter and enhancer, and calculated their flanking chromatin accessibility. To ensure reliability of analysis, we further filtered out TFs with unrobust score in promoter and enhancer as mentioned in methods, resulting in a selection of 100 qualified TFs from 134 TFs. Using consensus clustering, we separated the TFs into four clusters based on their change ratio of flanking chromatin accessibility on promoter or enhancer (Table S5). From Clusters 1 to Cluster 4, they represent the TFs, whose flanking chromatin accessibility in enhancer or promoter are highly resistant, moderately resistant, moderately sensitive and highly sensitive to BRM014 treatment (Fig. 6A). We found that some TFs has similar sensitivity to BRM014 at both enhancer and promoter, however some TFs display different sensitivity to BRM014 between at enhancer and promoter. Generally, flanking chromatin accessibility of TFs in enhancers is decreased more dramatically than in promoter, indicating that the sensitivity of TFs to BRM014 can be chromatin context-dependent, (Fig. 6B-6C).

**Figure 6:**
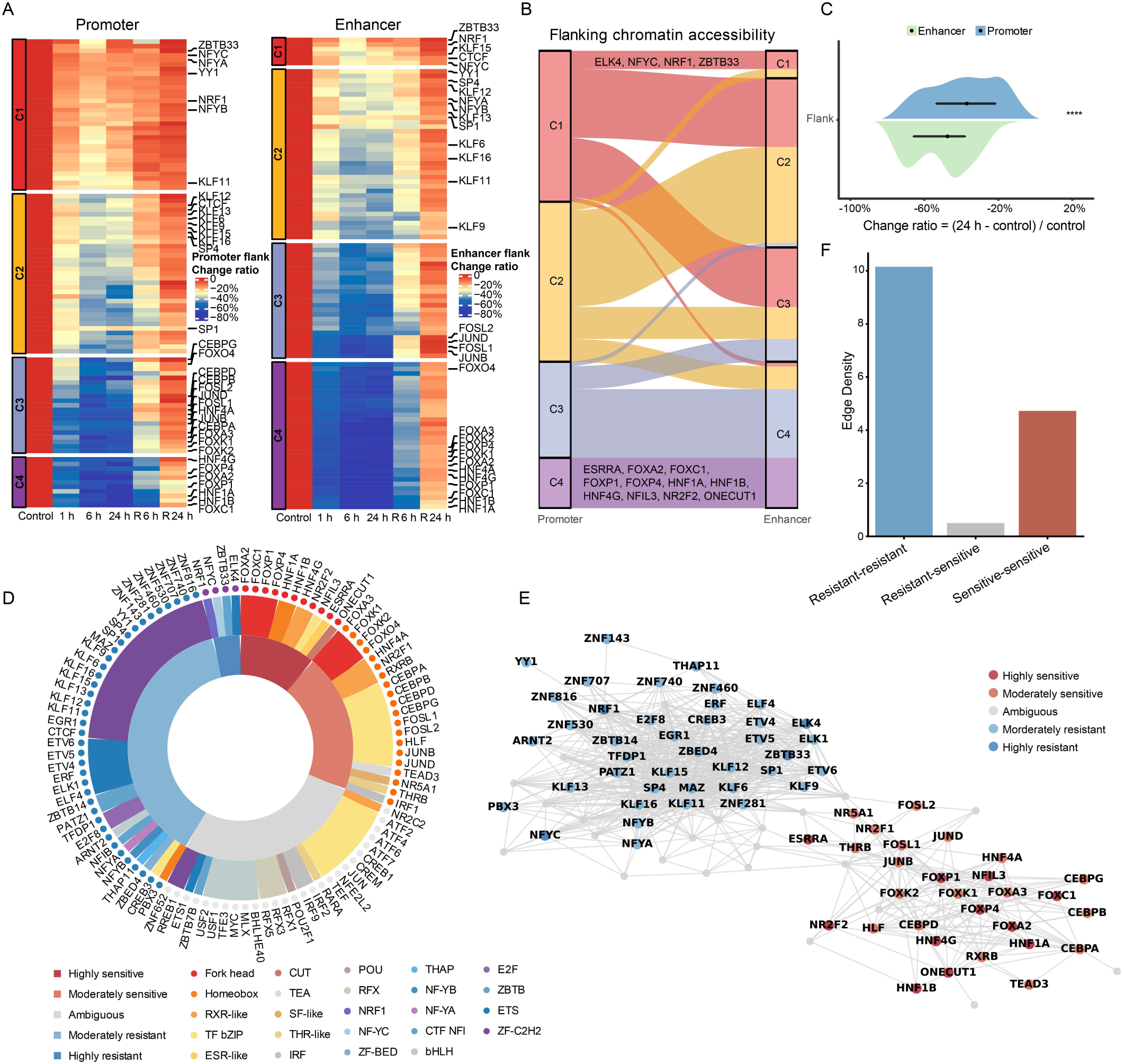
Definition of TF dependence on SWI/SNF reveals the spatial organization rule of TFs. **A.** Heatmap showing the change ratio of flanking chromatin accessibility around TF binding sites at promoter (left) and enhancer (right) for 100 TFs in HegG2 cells. TFs are grouped into four clusters based on their change ratios at different time points following BRM014 treatment. Grids color represents the dynamic change ratio compared with controls **B.** Sankey diagram illustrating the correspondence between the enhancer and promoter clustering categories for each TF as defined in panel A. TFs highlighted in cluster 1 or cluster 4 in both promoter and enhancer regions represent those that are highly resistant or highly sensitive to BRM014 treatment. **C.** Violin plot showing the distribution of change ratios in flanking chromatin accessibility for enhancers (green) and promoters (blue) in HepG2 cells after 24 h of BRM014 treatment, with statistical significance indicated **** (p.value <0.0001). **D.** Sunburst plot illustrating the sensitivity of TFs to BRM014 treatment, organized by clusters and families in HepG2 cells. The inner circle categorizes TFs into highly sensitive (dark red), moderately sensitive (light red), ambiguous (gray), moderately resistant (light blue), and highly resistant (dark blue) groups. The middle circle indicates TF family classifications, with each family color-coded as shown in the legend on the bottom. Each TF is labeled around the circular plot, with colored dots representing their respective categories of BRM014 sensitivity. **E.** The network diagram displays the Transcription factor co-occurrence network with the enhancer regions of HepG2. Each node represents a different TF, with the color coding indicating varying levels of sensitivity or resistance. Edges between nodes indicate potential co-occurrence relationships among the TFs **F.** The bar graph displays the edge density in transcription factor co-occurrence network within the enhancer regions of HepG2 cells among transcription factor pairs, categorized by their sensitivity to BRM014. The categories include interactions between TF pairs that are both resistant (blue bar), one resistant and one sensitive (grey bar), and both sensitive (red bar). See also Figures S9, Table S5

We defined the sensitive TF and resistant TF based on their sensitivity consistency of flanking chromatin accessibility on promoter and enhancer. The TFs are classified to highly resistant TF (C1 in both promoter and enhancer), moderately resistant TF (either C1or C2 in both promoter and enhancer), moderately sensitive TF (either C3 or C4 in both promoter and enhancer), and highly sensitive TF (C4 in both promoter and enhancer) (Fig. 6B). Resistant TFs enrich CTCF, NRF1, NFYA/B/C, YY1, ZBTB33, SP1/4, KLF proteins, ETS proteins and ZNF proteins, while sensitive TFs enrich HNF tissue specific factors, RXR-like family, FOX family, AP-1 family, and C/EBP family (Fig. 6D). TFs within the same family recognizing similar motifs shared comparable SWI/SNF dependency, highlighting evolutionary conservation in SWI/SNF dependency (Fig. 6D).

To characterize the functional differences of TFs in each cluster, we evaluated each TF’s regulatory preference by calculating the proportion of its binding sites overlapping promoter regions (±1 kb) of HepG2-specific versus housekeeping genes. Violin plot analysis showed that SWI/SNF-sensitive TFs are more likely to target cell-type–specific genes, whereas insensitive TFs preferentially bind to housekeeping gene promoters (Fig. S9B, left). GO enrichment analysis of the HepG2-specific gene set revealed that the top enriched terms were related to small-molecule catabolism and biosynthesis, consistent with the known metabolic functions of liver cells (Fig. S9B, right). Together, these results suggest that SWI/SNF-sensitive TFs are associated with specialized regulatory programs, while insensitive TFs are involved in fundamental cellular processes.

By integrating SWI/SNF dependency information, we further analyzed TF organizational patterns in enhancer and promoter in HepG2 cells by TF-COMB. Interestingly, TFs with the same SWI/SNF dependency prefer to locate closer to each other, irrespective of at enhancer and promoter (Fig. 6E and S9C), with co-occurrence of TFs within the same group significantly more than TFs between different dependency groups (Fig. 6F, S9D). Given that chromatin remodelers reshape chromatin by consuming ATP, we hypothesize that this spatial organization ensures both stability and plasticity of chromatin organization during genome regulation of development and signaling response in an energy-efficient manner. Further characterization of TF organization in other cells is crucial to determine if this spatial organization is conserved and its significance in gene regulation.

## Discussion

Genomic-wide capture of TF binding dynamics on chromatin is essential for unraveling gene regulatory networks and understanding cellular transcriptional changes during development and in response to external signals^3, 7^. We developed cFOOT-seq, which employs dsDNA deaminase to encode chromatin structure information into genomic DNA sequences. This approach enables simultaneous profiling of chromatin accessibility, nucleosome positioning, and TF occupancy on a genome-wide scale. cFOOT-seq measures TF occupancy based on chromatin footprints in both open and closed chromatin, capturing the dynamics of hundreds of TFs in a single experiment. Using known TF binding information and cFOOT-seq data, FootTrack quantitatively detects the dynamics of TF occupancy in transcriptional responses. FootTrack can also de novo predict TF occupancy from footprints and motifs, identifying potential TF candidates in transcriptional regulation. As a proof of concept, cFOOT-seq successfully detected TF dynamics during OKSM-mediated reprogramming, profiled TF dependencies on SWI/SNF, and uncovered a spatial organization rule where TFs with the same SWI/SNF dependence localize closely in HepG2. Thus, cFOOT-seq provide a novel genomics toolkit for studying gene regulatory networks and chromatin organization.

By encoding chromatin structure into DNA sequences, cFOOT-seq preserves DNA integrity, allowing for single-molecule and single-cell TF occupancy detection (Fig. 2H-I). When combined with ATAC-seq, cFOOT-seq enriches open chromatin regions, improving TF binding detection at lower costs. The increased sequencing depth in open chromatin enables more sensitive and quantitative TF footprint detection at single-molecular level. cFOOT-seq is compatible with long-read sequencing technologies like PacBio and Nanopore, as well as other chromatin probing methods such as DNase-seq, ChIP-seq, and CUT&Tag. Integrating plate-based single-cell technology with ATAC-cFOOT-seq (Fig. S5E-G, and 2I) enables TF occupancy detection in individual cells, offering potential insights into gene regulatory networks in heterogeneous populations and complex tissues.

Traditionally, genome-wide mapping of regulatory factor binding sites relied on antibody-based methods like ChIP-seq and newer techniques like CUT&RUN and CUT&TAG, which detect one protein at a time. In contrast, footprint-based methods can simultaneously address all factors in one experiment, identifying occupied sites with nucleotide precision. cFOOT-seq data offers comprehensive, high-resolution insights into the spatial and temporal dynamics of TF binding, elucidating regulatory networks and gene expression mechanisms in various biological processes.

### Limitations

Despite its strengths, cFOOT-seq has certain limitations, including residual enzyme sequence bias, high sequencing costs, and challenges in transcription factor (TF) assignment. SsdAtox exhibits lower sequence bias than DddA, Ddd_Fa, and Ddd_Ss^48^. DddB was recently applied in the FOODie method for footprint detection; however, the relative bias differences between DddB and SsdAtox remain unclear and have not been systematically compared^80^. Current correction strategies mitigate much of the bias, but residual effects persist, particularly for certain motifs. Further improvements, such as deep learning-based correction models^81, 82^, may enhance analytical accuracy. Additionally, engineering SsdAtox variants with reduced sequence bias could further improve the method.

Reliable genome-wide TF footprint detection by cFOOT-seq requires approximately 100M reads per sample, with biological replicates reducing random noise. Combining cFOOT-seq with ATAC-seq can reduce sequencing costs and enhance detection. However, each combined approach has its own advantages and limitations. ATAC-cFOOT-seq, which integrate Tn5-mediated open chromatin enrichment before deamination, provides detection of TF footprint with significant reduced sequencing depth, but may miss some TF footprints due to the Tn5 incubation steps. In contrast, cFOOT-ATAC-seq, while offering lower open chromatin enrichment, allows for sensitive TF footprint detection.

The lack of known motifs for certain TFs limits the scope of cFOOT-seq. We mainly used motifs from JASPAR, and incorporating more databases could expand TF coverage. Some TFs with defined motifs still show weak or no footprints, possibly due to dynamic binding or technical loss during the experimental process. Moreover, similar motifs shared by different TFs make footprint assignment challenging, a common issue for footprint-based methods. Expanding the motif repertoire to include novel motifs for uncharacterized TFs, as well as composite motifs of TF pairs, would further improve the applicability of cFOOT-seq. Enhanced understanding of TF binding grammar, including DNA sequence, 3D shape, and neighboring motifs, would improve TF binding prediction accuracy.

## Acknowledgements

This work was supported by the National Key Research & Developmental Program of China (2021YFA1102000, 2022YFA1106000, 2021YFA1102900), the National Natural Science Foundation of China (No.32270775, No.81902583, No.32270702, No.32370868, No. 32070802), the Fundamental Research Funds for the Central Universities (No. 22120250374), and Peak Disciplines (Type IV) of Institutions of Higher Learning in Shanghai. We thank Dr. Yangming Wang (Peking University) for generously providing plasmid constructs, and Dr. Meng Qu (Zhejiang University) for kindly sharing cell lines used in this study. We also thank members of the Zhang, Gao, Shi and Liu laboratories for their helpful suggestions and discussions on the subject over the course of the study.

## Author Contributions

J.-M.Z., S.G., J.S., and X.L. designed the study and guided the project. H.W., and M.-C.Y. performed majority of the experiments. A.W. analyzed majority of the data. Z.D. performed the scATAC-cFOOT-seq. Z.S., K.C., B. H., Y.F., X.W., X.-F.C., B.H., Y.F. performed part of the experiments. X.C., Y.Z., Y.-X. L. analyzed part of the data. L.K., Y.H., K.C., S.B., J.T., J.X. and C.W. provide key reagents and important suggestions.

## Supplemental information

Document S1: Figures S1-S9

Table S1-S8

Table S1. reference_data_source

Accession information for various TF ChIP-seq, ATAC-seq, and other datasets across different cell lines (HepG2, K562, R1, MEF).

Table S2. scATAC-cFOOT-seq

The barcodes used for single cell library preparation and different metrics of single cell data in scATAC-cFOOT-seq

Table S3. chomHMM_cluster

Group information of 204 TFs in HepG2 cells clustered by their TFOS in regions defined by chromHMM annotations

Table S4. BRM014_TFOS and Flank_change ratio_all sites

Clustering information of 134 TFs based on their dynamic responses to BRM014 treatment, with change ratio of TFOS and Flanking chromatin accessibility at all binding sites of each TFs in HepG2

Table S5. BRM014_Flank change ratio_ Promoter_Enhancer

Clustering information of 100 TFs in HepG2 based on their change ratio of flanking chromatin accessibility on promoter or enhancer.

Table S6. Oligo sequence data

The sequence of oligos used in this study Table S7. human_mouse_cluster

Clustering of TFs in human and mouse based on the similarity of binding motifs. Table S8. cFOOT_data

The cFOOT-seq sequencing data generated in this study are listed.

## Methods

### Lead contact and materials availability

Further information and requests for reagents may be directed to the Lead Contact, Jia-Min Zhang.

### Cell culture

All cells were cultured according to standard procedures.

HepG2 cells (Procell) were maintained in MEM with NEAA (Procell) and K562 cells (Procell) were maintained in RPMI 1640 (Thermo Fisher Scientific).

Mouse embryonic fibroblasts (MEFs) were derived from 13.5 days mouse embryos (ICR), then maintained in DMEM-high glucose (Life Technologies) supplemented with 10% fetal bovine serum (FBS, Gibco), 1 mmol/L L-glutamine (Thermo Fisher Scientific). Puromycin-resistant MEF cells were derived from 13.5 days mouse embryos (DBA and C57 background), and puromycin-resistant feeder were obtained by treating these MEFs with mitomycin C for 3 h.

mESC medium including DMEM-high glucose medium, supplemented with 15% FBS, 1 mmol/L L-glutamine, 100× nucleosides (EMD Millipore), 100× NEAA (Millipore), 0.11 mmol/L 2-mercaptoethanol (Sigma-Aldrich), 10^3^ U/mL LIF (Millipore) and 100× penicillin/streptomycin (Gibco). mESC were cultured on feeder, which was obtained by treating MEF cells (ICR) with mitomycin C for 3 h.

The mESC cell line R1 (includes wild-type and knockout cell lines) were cultured in mESC medium supplemented with 1 μmol/L PD0325901 (Selleck) and 3 μmol/L CHIR99021 (Selleck), while mESC-OG2 cell line used in BRM014 treatment experiment maintained in mESC medium. The OG2 cell line is previously generated in Gao lab^83^.

### Cell line generation

We established rtTA-Cas9-EGFP mESC lines and selected three Yy1 sgRNA sequences, which were cloned into the lentiGuide vector (Addgene). Each sgRNA-containing vector (Table S6) included markers for BFP, RFP, and puromycin resistance, respectively. The lentiGuide-Yy1 sgRNA plasmids were extracted and purified using the EndoFree Plasmid Maxi Kit (CoWin Biotech Co.).

To produce lentivirus, the vectors were transfected into 293T cells along with packaging plasmids psPAX2 and pMD2G. Harvested lentivirus after transfection 72 h, then infected 3×10^4^ rtTA-Cas9-EGFP mESCs with three types lentiGuide viruses simultaneously. After 8-10 h of virus infection, cells were washed with culture medium to remove residual virus and then maintained in fresh medium.

Following lentiviral infection, cells were selected with puromycin for 7 days, during the selection, mESC were cultured on the puromycin resistant feeder. BFP and RFP-positive mESC were sorted using the CytoFLEX SRT Cell Sorter (Beckman Coulter). Subsequently, Cas9 protein expression induced by doxycycline and Yy1 knockout efficiency was confirmed by Western blot.

### Purification of SsdA_tox_, Ddd_Ss, and Ddd_Fa for deamination assay

The expression and purification of SsdA_tox_ were performed as previously described, with minor modifications^47^. The pETduet-SsdA_tox_-SsdAI vector was introduced into Rosetta™ 2 (DE3) Singles™ Competent Cells (EMD Millipore). After overnight culture, transformed cells were inoculated into 4 liters of 2YT medium (1:100 dilution) and grown to approximately OD600 = 0.6. Protein expression was induced with 0.5 mM IPTG (Sigma-Aldrich), followed by incubation at 16°C for 18 h. Cells were harvested by centrifugation (4,000 rpm, 10 min) and resuspended in lysis buffer (50 mM Tris-HCl pH 7.4, 500 mM NaCl, 10 mM imidazole, 5 mM beta-mercaptoethanol, 100 uM PMSF, 1× protease inhibitor cocktail). Lysis was performed using a high-pressure cell cracker (JN-miniPro), and the lysate was clarified by centrifugation (28,000 g, 30 min). The his-tagged SsdA_tox_-SsdAI complex in the supernatant was purified using Ni-NTA beads (QIAGEN) by incubation at 4°C for 3 h. Then beads were washed by lysis buffer 3 times before the denaturation. To separate SsdA_tox_ from SsdAI, the beads underwent denaturation in denaturation containing 8 M urea for 16 h at 4°C, followed by renaturation through successive washes with lysis buffer containing decreasing urea concentrations (6 M, 4 M, 2 M, 1 M, 0 M). The refolded proteins bound to Ni-NTA beads were eluted by Elution buffer (50mM Tris-HCl pH7.4, 500mM NaCl, 300mM imidazole, 1mM DTT, 5% glycerol). Imidazole was subsequently removed by sequentially dialysis against dialysis buffer containing 500 mM NaCl (50 mM Tris-HCl pH 7.4, 500 mM NaCl, 1 mM DTT, 10% glycerol) and dialysis buffer containing 200 mM NaCl. Purity and quantification was assessed by SDS-PAGE, and high-quality fractions were stored at -80°C.

The expression and purification of Ddd_Fa and Ddd_Ss were performed as previously described^48^, with minor modifications. The pCOLADuet1-10His-Ddd_Fa-com, pCOLADuet1-10His-Ddd_Ss-com vectors (gift from Yangmin Wang’s lab in Peking Univerisity) were introduced into Rosetta™ 2 (DE3) Singles™ Competent Cells (EMD Millipore). The steps for culture, collection, lysis and protein bound with Ni-NTA beads are the same as purification of SsdA_tox_. To separate Ddd_Fa and Ddd_Ss from its immunity protein, the beads underwent denaturation in lysis buffer containing 6M GuHCl for 3 h at 4°C, followed by renaturation through successive washes by lysis buffer containing 10uM ZnCl_2_ with decreasing GuHCl concentrations (6 M, 5 M, 4 M, 3 M, 2 M, 1 M, 0M). The steps of elution, dialysis and quantification is the same as the protocol of SsdA_tox_.

### Deamination assay to analyze the deaminase activity on oligo and naked genomic DNA

DNA deamination assays on oligonucleotides were performed as previously described^46, 47^, with some modifications. All DNA substrate (Table S6) were purchased from Sangon Biotech (Shanghai, China) and contained a 6-FAM fluorophore at the 5′ end for visualization. To generate double-stranded (ds) DNA substrate, 6-FAM-labeled oligonucleotides were annealed with their complementary unmodified strands at a final concentration of 10 μM in 1x annealing buffer (10 mM Tris pH 7.5, 50 mM NaCl, 1 mM EDTA). Deamination reactions were carried out in a 10 μL mixture consisting of 10 mM Tris-HCl pH 7.4, 0.1 μM substrate, and deaminase at the concentrations indicated in Fig. S1. The reactions were incubated at 37°C for 1 hour. To terminate the deamination reaction, samples were heated at 85°C for 15 min, followed by the addition of 0.3 μL UDG (NEB) and further incubation at 37°C for 30 min. Substrate cleavage was induced by adding 0.5 μL of 2M NaOH and incubating at 95°C for 10 min. The mixture was then cooled on ice, combined with 10 μL of formamide and 2 μL of 6× purple DNA loading dye (NEB), and analyzed by 15% 8M urea-denaturing acrylamide gel electrophoresis in 0.5× TBE buffer. The 6-FAM signal was detected and quantified using the ChemiDoc MP imaging system (Bio-Rad).

For deaminase activity test on naked genomic DNA, reactions were performed in 20 μl deamination mixture containing 10 mM Tris-HCl pH 7.4, 1× PIC, 1 mM DTT, 0.8 ng lambda DNA, 0.1 U/μl Uracil Glycosylase Inhibitor (UGI, NEB), 80 ng genome DNA extracted from R1 cells, and deaminase at the concentrations indicated in Fig. S1. Reactions were incubated for 10 min at 37°C. DNA was precipitated by adding 20 μL of isopropanol and 2 μL of 3M NaAc, followed by incubation at -80°C for 1 h. The DNA pellet was then resuspended in 20 μL of ddH₂O. 10 ul DNA were used for genome-wide library preparation and sequencing, following the cFOOT-seq protocol.

### UPLC-MS/MS analysis for testing deaminase activity on DNA with C and 5mC

DNA substrate containing cytosine (C-DNA) or 5-methylcytosine (5mC-DNA) were prepared via PCR amplification using either dCTP or 5-methyl-dCTP (5m-dCTP) to replace dCTP. The PCR was performed with primers C-DNA/5mC-DNA_F and C-DNA/5mC-DNA_R, using C-DNA/5mC-DNA_template as the template and TaKaRa Taq™ Hot Start Version (TaKaRa) for amplification. The PCR product was purified using the SanPrep Column DNA Gel Extraction Kit (Sangon), following the manufacturer’s instructions. The sequences of the primers and template are described in Table S6.

For the deamination test, 40 ng of PCR-amplified DNA substrate was incubated with increasing concentrations of deaminase (0, 200, 400, 800, 1600, 3200 nM) in a 10 μL reaction mixture containing 10 mM Tris-HCl, 1× PIC, 1 mM DTT, and 0.1 U/μl UGI. The reaction was incubated at 37°C for 10 min with continuous mixing at 1000 RPM. After deamination, DNA was precipitated by adding 10 μL of isopropanol and 1 μL of 3 M NaAc, followed by incubation at -80°C for 1 h. The DNA pellet was then resuspended in 20 μL of ddH₂O.

To measure the deamination efficiency, the purified deaminated DNA was first digested into single nucleosides by nuclease P1 (NEB) and then dephosphorylated by Quick CIP (NEB) according to the manufacture’s instructions. The nucleoside composition of the DNA samples were detected by ACQUITY UPLC system (Waters) and Triple Quad™ 6500+ LC-MS/MS system (SCIEX) using multiple reaction monitoring (MRM) mode with an ACQUITY Premier HSS T3 (100 Å, 1.8 μm, 2.1 × 100 mm, Waters) column. The flow rate was set at 0.3 mL/min with mobile phases A (water with 0.1% formic acid) and B (100% methanol). The linear gradient was as follows: 100% to 95% A (0-2 min), 95% to 90% A (2-4 min), 90% to 50% A (4-7 min), 50% to 5% A (7-7.5 min), 5% A (7.5-9 min), 5% to 100% A (9-9.1 min) and 100% A (9.1-10 min). The quantifier transitions for each nucleoside were: dC: 228.1/112.1 (CE 20, DP 20); dG: 268.1/152.1 (CE 20, DP 60); 5mC: /126.1 (CE 17, DP 20). And retention time for each compound were: 5mC: 3.75 min; dC: 2.38 min; dG: 5.05 min. All compounds were measured in positive ESI mode and were quantified by interpolating the peak areas of the quantifier MRM transitions from the standard curves.

### cFOOT-seq procedures

#### Nuclei preparation

Collected 5×10^4^ cells for each sample, washed with cold DPBS-0.04% bovine serum albumin (BSA), centrifuge at 300g, 4°C, 5min, remove supernatant. The pellets were resuspended in 50 μl cold nuclei extraction buffer (modified Omni-ATAC buffer: 10 mM Tris-HCl, 10 mM NaCl, 3 mM MgCl2, 0.1% Tween-20, 0.1% IGEPAL CA-630, 0.01% Digitonin, 0.1 mM EDTA, 1× PIC, 1%(w/v) BSA). Incubate on ice 3min, then add 1ml cold stop buffer (10 mM Tris-HCl, pH 7.4, 10 mM NaCl, 0.1mM EDTA, 1×PIC, 1%(w/v) BSA) and invert tube 3 times gently mix. The nuclei were collected through centrifugation at 700g, 4°C, 10 min. Carefully remove the liquid above without disturbing the sediment.

#### Deaminase Reaction

Resuspend the pellet gently in 50 μl deamination mix (10 mM Tris-HCl, 1× PIC, 1 mM DTT, 2 ng lambda DNA, 0.1 U/μl UGI, 5-15U/μl SsdA_tox_), incubate at 37°C for 10 min with shaking at 1000 RPM. The concentration of SsdA_tox_ required for each cell lines should be tested to generate general genomic conversion rate between 25-40%. Longer incubation increases the conversion rate but may compromise footprint detection due to potential loss of TF occupancy on chromatin.

#### DNA Extraction

After the reaction, add 10% SDS and 1 M NaHCO_3_ to final 250 μl with 1% SDS and 0.1M NaHCO_3_, and incubate at 65°C for 1.5h with shaking at 1350 RPM. Then add 250 μl PCI and mix by full-speed vortexing for 5s. Remove the upper liquid to a new tube and then precipitate DNA with equal volume isopropanol.

#### Genome-Wide Library Preparation and Sequencing

Fragment genomic DNA using Covaris DNA shearing (peak power: 50.0, duty factor: 20.0, cycles/burst: 200, duration: 25 sec). Prepare the DNA library using the EpiArt DNA Methylation Library Kit for Illumina V3 (Vazyme) as described. Briefly, 20 ng of input genomic DNA was denatured to single-stranded DNA, followed by the ligation of a truncated linker at the 3’ end. The single-stranded DNA was then extended to double-stranded DNA using extension primers. After purification with 1.2× VAHTS DNA Clean Beads (Vazyme, N411), a truncated adapter was ligated to the 5’ end of the original template strand of the double-stranded DNA. The reaction products were further purified with 1× VAHTS DNA Clean Beads, and the complete library was amplified with universal i5 and i7 primers for 7 cycles. Then the complete library was purified by 0.85× VAHTS DNA Clean Beads, and eluted by 10-20ul elution buffer.

Finally, paired-end 150 bp sequencing was performed on the Illumina NovaSeq 6000 system. For condition test, approximately 7 million read pairs per sample (∼1.5× depth) are sufficient. For de novo footprint prediction, approximately 100 million read pairs per sample (∼6.5× depth) are recommended for downstream analyses. More details can be found in Table S8.

### cFOOT-seq procedures for small number of cells with ConA beads

#### Nuclei preparation

Collect the appropriate number of cells, wash with 500 μl wash buffer (20 mM HEPES, 150 mM NaCl, 0.5 mM Spermidine, 0.1% BSA, 1 × PIC). For cell counts between 1000 and 5000, dilute as needed, and for 5 to 200 cells, a stereomicroscope may be used. Resuspend the cells with 300 μl binding buffer (20 mM HEPES-KOH, 10 mM KCl, 1 mM CaCl_2_, 1 mM MnCl_2_). After resuspending 5 μl of Concanavalin A-coated magnetic beads with 15 μl binding buffer, mix with the cell suspension and incubate for 10 min, place the EP tube on a magnetic stand for 1 min, discard the liquid and remove the EP tube, then use 500 μl of cold blocking buffer (20 mM HEPES, 150 mM NaCl, 0.5 mM Spermidine, 0.1% BSA, 2 mM EDTA, 1× PIC, 0.01% digitonin) to resuspend the beads and incubate for 5 min. Repeat the magnetic separation and liquid removal. Subsequently, resuspend the beads in 500 μl of cold dig-Wash Buffer (20 mM HEPES, 150 mM NaCl, 0.5 mM Spermidine, 0.1% BSA, 1× PIC, 0.01% digitonin) and wash twice more using the same procedure.

#### Deaminase Reaction and DNA Extraction

Resuspend the beads with 100 μl reaction buffer (10 mM Tris-HCl pH=7.4, 1× PIC, 1 mM DTT, 2 ng lambda DNA, 0.1 U/μl UGI, 2-10 U/μl SsdA_tox_), incubate at 37 °C, shaking at 1000 rpm for 10 min. Add 12.5 μl of NaHCO_3_ and 12.5 μl of 10% SDS, mix and incubate at 65 °C, shaking at 1350 rpm for 1 h. Place the EP tube on the magnetic stand for 1min, and transfer the liquid to a new EP tube. Add 125 μl of PCI and vortex at full speed for 5 seconds to mix. Carefully transfer the upper liquid to a new tube and precipitate the DNA with an equal volume of isopropanol.

#### Genome-Wide Library Preparation and Sequencing

The procedure closely follows that of cFOOT-seq, with slight modifications. Given the low quantity of genomic DNA, the entire sample is used for fragmentation. The fragmented DNA is then enriched using VAHTS DNA Clean Beads beads before denaturation and subsequent ligation of the 3’ adapter with the EpiArt DNA Methylation Library Kit for Illumina V3 (Vazyme).

### cFOOT and ATAC-seq combination procedures

There are two protocols included ATAC-cFOOT-seq and cFOOT-ATAC-seq. Two shared steps of Tn5 transposase generation, and DNA library preparation steps are done as below.

#### Tn5 Transposome Generation with Cytosine-Free Adaptor

To avoid cytosine deamination of Tn5 adapters by SsdA_tox_, a cytosine-free Tn5 adapter was prepared by annealing Tn5 Primer D (Table S6), which lacks cytosine in its 5’ overhang, with the universal Tn5 ME oligo (5‘-phos-CTGTCTCTTATACACATCT-NH2-3’). The annealing was performed in a thermal cycler with an initial denaturation at 95°C for 5 minutes, followed by a gradual cooling to 10°C at a rate of -0.1°C/s. Subsequently, 5 µM of the cytosine-free adaptors were incubated with unloaded Tn5 transposase protein in coupling buffer (Vazyme, S601) at 30°C for 1 hour, producing a final concentration of 750 nM Tn5 transposome. The loaded Tn5 transposase can be stored at -20°C for up to 1 year.

#### DNA library preparation for combined procedures

DNA library was constructed with modification of protocols used in the EpiArt DNA Methylation Library Kit for Illumina V3 (Vazyme). In brief, denature 50 ng genomic DNA, ligate a truncated linker at the 3’ end, extend to full duplex, and purify with 1.2 x VAHTS DNA Clean Beads. Then purified DNA is initially amplified for 4 cycles with universal i7 primers and i5_bridge primer (Table S6) to add 5’ sequencing fragment. After purification by 1× VAHTS DNA Clean Beads, the library is further amplified with universal i5 and i7 primers for 8 cycles, which can match illumine next generation sequencing platform. Finally, the library fragments in the size range of 300-500 bp can be recovered by agarose gel electrophoresis to obtain the final library.

### ATAC-cFOOT-seq protocol

#### Nuclei preparation

The operation is same as that in cFOOT-seq.

#### Tn5 transposase reaction

Resuspended nuclei pellet gently with 50 μl tagmentation mix (10mM Tris-HCl pH=7.6, 5mM MgCl_2_, 10% Dimethylformamide (DMF), 33%(v/v) DPBS, 100nM Tn5 transposome with cytosine-free adaptor), incubate reaction at 37°C for 15min in a mixer with 1000RPM.

#### Deaminase Reaction

Stop Tn5 reaction by adding 500 μl cold tagmentation stop buffer (10mM Tris-HCl pH=7.4, 1mM DTT, 1x PIC), mix well, collect nuclei through centrifugation at 700g, 4°C, 5min. Remove all supernatant, resuspension pellet by deamination mix (10mM Tris-HCl pH=7.4, 1× PIC, 1mM DTT, 0.2ng lambda DNA with Tn5 adaptor sequence, 0.1 U/μl UGI, 7.5-15U/μl SsdA_tox_), incubate reaction at 37°C for 10 min in a mixer with 1000 RPM.

#### DNA Extraction and library preparation

The operation of DNA extraction is same as that in cFOOT-seq. DNA library is prepared as described in DNA library preparation for combined procedures.

#### Sequencing

paired-end 150 bp sequencing was performed on the Illumina NovaSeq 6000 system. For condition testing, approximately 3 million read pairs per sample (∼6× depth in open chromatin regions) are sufficient. For de novo footprint prediction, approximately 60 million read pairs per sample (∼42× depth in open chromatin regions) are recommended.

### cFOOT-ATAC-seq protocol

#### Nuclei preparation

The operation is same as that in cFOOT-seq.

#### Deaminase reaction

Resuspended nuclei pellet by 50 μl deamination mix (10mM Tris-HCl pH=7.4, 1× PIC, 1mM DTT, 33%(v/v) DPBS, 0.02% BSA, 0.1 U/μl UGI, 1-5U/μl SsdA_tox_), incubate reaction at 37°C for 10 min in a mixer with 1000 RPM.

#### Tn5 transposase reaction

Stop deaminase reaction by adding 500 μl cold deamination stop buffer (10mM Tris-HCl pH=7.4, 1× PIC, 1mM DTT, 33%(v/v) DPBS, 0.02% BSA, 5mM MgCl_2_). Collect nuclei through centrifugation at 700g, 4°C, 10 min. Remove supernatant, resuspension pellet by 50 μl tagmentation mix (10mM Tris-HCl pH=7.6, 5mM MgCl_2_, 10% Dimethylformamide (DMF), 33%(v/v) DPBS, 100nM Tn5 transposome with cytosine-free adaptor). Incubate reaction at 37°C for 15-30 min in a mixer with 1000 RPM.

#### DNA Extraction and library preparation

The operation of DNA extraction is same as that in cFOOT-seq. DNA library is prepared as described in DNA library preparation for combined procedures.

#### Sequencing

paired-end 150 bp sequencing was performed on the Illumina NovaSeq 6000 system. For condition testing, approximately 3 million read pairs per sample (∼2× depth in open chromatin regions) are sufficient. For de novo footprint prediction, approximately 60 million read pairs per sample (∼13× depth in open chromatin regions) are recommended.

### Single cell ATAC-cFOOT-seq procedures

#### Nuclei preparation and deaminase reaction

Follow the same steps as in ATAC-cFOOT-seq.

#### DAPI staining and FACS sorting

After the deaminase reaction, collect nuclei through centrifugation at 700 g, 4°C, 5 min. Remove all supernatant, resuspend the pellet in DPBS-0.5% BSA containing DAPI (1:1000) and incubate in the dark on ice for 15 min. After incubation, centrifuge and remove all supernatant, resuspend the nuclei in DPBS-0.5% BSA without DAPI to prepare for sorting. Then sort the DAPI-positive single nuclei into single wells of a 96-well plate containing 1μl lysis buffer (10 mM Tris pH 8.0, 20 mM NaCl, 1 mM EDTA, 0.1% SDS, 500 nM Carrier ssDNA, 60 μg/mL protease K (QIAGEN)) by CytoFLEX SRT Cell Sorter (Beckman Coulter). After sorting, immediately seal the single-cell lysis plate and centrifuge at 1,000 g for 1 min at RT to avoid losing the cells on the plate wall.

#### Nuclei lysis and Tn5 release

After centrifugation, incubate the single-cell lysis plate at 65 °C for 15 min, 95°C 2 min in a thermal cycler with a heated lid set to 80 °C. Then add 1 μl 3% Triton X-100 to quench SDS and stop the lysis reaction. After this, store the plate at -80 °C or proceed to downstream amplification.

#### Library preparation

The DNA library was constructed following a modified protocol based on the EpiArt DNA Methylation Library Kit for Illumina V3 (Vazyme). Briefly, single-cell genomic DNA was denatured in the plate, and a truncated linker was ligated to the 3’ end. The ligated DNA was amplified for 8 cycles using universal i7 primers and i5_bridge primers with well-specific barcodes to distinguish single cells in each well. The amplified products from all wells were pooled and purified twice using 1× magnetic beads. The library was then further amplified for 12 cycles with universal i5 and i7 primers to ensure compatibility with the Illumina next-generation sequencing platform and purified using 0.85× magnetic beads. To achieve a total DNA yield of 300-500 ng, an additional 6 cycles of amplification are required, followed by purification using 0.85× beads. Finally, library fragments in the size range of 300-500 bp were recovered by agarose gel electrophoresis to obtain the final library.

#### Sequencing

paired-end 150 bp sequencing was performed on the Illumina NovaSeq 6000 system, with approximately 2 million read pairs per cell. More details can be found in Table S2.

### OSKM reprogramming

The reprogrammable MEF cell line was generated in previous study ^84^, doxycycline was added to induce MEF cells overexpress OSKM at a final concentration of 1 µg/ml. Samples were collected at 24, 48, and 96 h after doxycycline induction.

### BRM014 treatment

BRM014 compounds (BRM/BRG1 ATP Inhibitor-1, MedChem Express) dissolved in dimethyl sulfoxide (DMSO) to final concentration of 10 mM. Add the required concentration directly to the well and shaken thoroughly.

For the recovery, the medium was used to gently flush the well 3 times and finally to medium without BRM014.

### cFOOT-seq reads preprocessing

The raw sequencing reads obtained from next-generation sequencing were first evaluated for quality using FastQC v0.12.1. Subsequent trimming of these reads was performed with Trim Galore v0.6.10, employing specific parameters to enhance both quality and utility for downstream analysis. These parameters included --trim-n, --clip_R1 3, --clip_R2 10, --three_prime_clip_R1 3, -- three_prime_clip_R2 3, --length 35, -q 20, --fastqc, --paired. The alignment of the trimmed reads was conducted using BASAL (https://github.com/JiejunShi/BASAL)^85^, which is specifically designed for mapping nucleotide base-conversion sequencing reads in a conversion-sensitive manner. The alignment parameters were set to -m 1, -x 1000, -r 1, -v 0.06, -s 16, -S 1, -n 0, -g 1, - M C:T. For further processing, the BasalKit^85^ ‘avgmod’ module was utilized with parameters -r -m 1 -i correct to process the aligned reads, producing a TSV file that recorded the average C-to-T conversion rate for each cytosine. In refining the analysis, SNP sites were systematically excluded. For detailed analysis of TF occupancy, the study proceeded with two analytical approaches: analysis based on known transcription factor binding sites and de novo prediction of transcription factor binding sites.

### FootTrack analysis

To comprehensively assess transcription factor occupancy and dynamics across variable conditions, we developed the FootTrack (Footprint Analysis for Tracking TF Occupancy and Kinetics). Constructed on the basis of TOBIAS^37^, FootTrack serves as an extensive analytical platform tailored for both the analysis of known transcription factor binding sites and the prediction of new sites. The tool’s structured analytical process comprises three main components: bias correction, analysis of known transcription factor binding sites, and de novo prediction of transcription factor binding sites. Together, these elements enable a robust examination of transcription factor interactions within the genomic landscape.

#### 1. Bias correction

To assess enzymatic bias, data from naked DNA samples with a conversion rate of approximately 40% were utilized, as this rate closely aligns with the conversion rate observed in the open chromatin regions of cFOOT-seq. Bias was quantified by aggregating conversion rate across different sequence contexts and normalizing them by the frequency of each context in the background. A position-weight matrix (PWM) was generated using the sequence context within a ±10 bp window, and this PWM was subsequently applied in further analyses to compute bias for each context.

For the correction process, FootTrack first calculates the background conversion rate based on the observed data. This background rate is then multiplied by the context bias to obtain the expected conversion rate for each base. The corrected conversion rate is determined by subtracting the expected conversion rate from the observed value^37^.

Two background calculation modes are available. The first, the global background, uses the average conversion rate across the entire genome, preserving both chromatin accessibility and transcription factor binding information. The second, the local background, calculates the average conversion rate within a ±50 bp window around each base^86^. This mode improves TF footprint detection by reducing biases from regional chromatin accessibility variations. The correction formula is as follows:

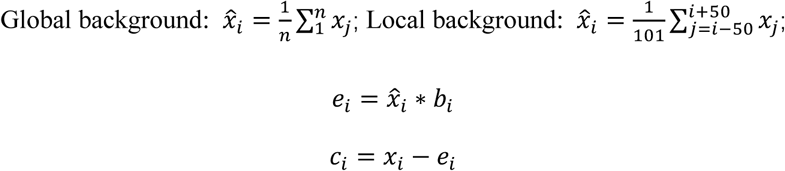

Where:

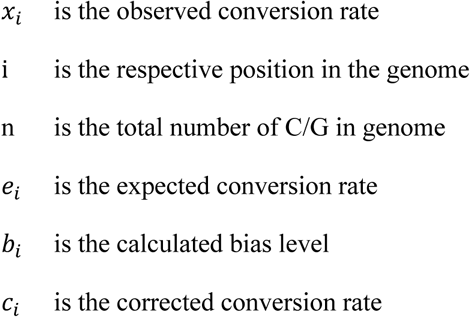

The corrected conversion rate are then used in subsequent analyses to improve the accuracy of transcription factor binding site identification and characterization.

#### 2. Analysis based on known transcription factor binding sites

Potential TF binding sites are identified by scanning motifs in the peak regions defined by ChIP-seq, with consideration of strand orientation. The flanking regions are defined as extending 50 bp on either side of each motif. The FOS at each site is calculated by subtracting the conversion rate in the flanking regions from that at the motif site. Averaging the FOS across all binding sites or specific regions gives the Transcription Factor Occupancy Score (TFOS), which reflects overall TF occupancy. The data sources for TF binding information are provided in Table S1.

#### 3. Analysis based on de novo prediction of transcription factor binding sites

FootTrack provide two strategies for footprint detection and motif prediction:

1. For cFOOT-seq data, we propose strategy 1 (S1), adopted from TOBIAS^37^. S1 calculates footprint scores for each motif with the corrected data, and applies statistical methods to identify motifs with higher scores as potential TF binding sites in interested regions. *Footprint Scoring calculation*: FootTrack employs footprint scoring to predict TF binding sites by assessing the likelihood of TF binding on genome. The footprint score is computed by analyzing the mean conversion rate of two strategically defined genomic regions: the ‘center’ and the ‘flanks.’ Specifically, the ‘center’ is defined by the interval [i + w_*_, i + w_*_ + w_+_], with w_*_, the flank width, set by default to 20 nucleotides, and w_+_, the center width, to 10 nucleotides. The ‘flanks’ are the regions extending from i to i + w_*_ and from i + w_*_ + w_+_ to i + 2 w_*_ + w_+_. The footprint score is determined by subtracting the mean conversion rate of the ‘center’ from that of the ‘flanks’. *TF Binding Site Detection*: FootTrack uses the MOODS^87^ (https://github.com/jhkorhonen/MOODS) to scan for motifs, applying a p-value threshold of 1e-4. For each identified TF motif, footprint scores are calculated and matched accordingly. Background base pair probabilities are estimated based on the open chromatin regions, and a background distribution of scores is generated by randomly subsetting these regions at ∼200 bp intervals. Each TF motif is then classified into bound and unbound sites based on a score threshold. This threshold is determined by the significance level derived from a normal distribution fit to the background score distribution, with a p-value cutoff of 0.05. Open regions are usually used for de novo prediction, considering the enrichment of TF binding.
2. For ATAC-cFOOT-seq and cFOOT-ATAC-seq, we proposed strategy 2 (S2), conceptionally adopted from Footprint tools^40^. S2 initially detects footprint regions by binomial statistical test and then scans for motifs within the identified footprint regions to predict TF binding sites. *Footprint Detection*: The BasalKit^85^ ‘avgmod’ module is used to extract data from the mapped BAM files, providing the number of reads that cover each base and the number of converted reads at each base. The expected conversion rate for each base is derived from the previous bias correction step. The number of converted reads at each base is treated as the number of successes in a series of independent trials, allowing the observed data to be modeled using a binomial distribution. By comparing the observed data to the expected data, we assess whether the observed conversion rate is significantly lower than expected. Specifically, the null hypothesis (H_0_) posits that the observed conversion rate at each base equals the expected. The alternative hypothesis (H_1_) suggests that the observed conversion rate is significantly lower than expected, indicating potential protein binding at the corresponding position. In our analysis, a threshold of p<0.05 is used to classify a position as likely to be a footprint. Since deamination occurs only at C/G sites, and to minimize the proportion of missing values, we extend the footprint by 5 bp on either side to define the final footprint region. *TF Assignment*: Similar to the approach used in S1, once TF motifs within open chromatin regions are identified, any motif that overlaps with a footprint by more than 50% is considered a potential transcription factor binding site

#### 4. Differential transcription factor binding analysis

FootTrack employs volcano plots to visualize differences in TF occupancy between different cells and conditions. TFs are selected based on significant differential footprint scores greater than 0.1 and a -log10 (p-value) exceeding the 90th percentile, identifying them as condition-specific TFs. To minimize false positives due to motif similarity, the TOBIAS ClusterMotifs tool is used to group motifs by similarity, with a threshold of 0.4 for robust clustering. Each cluster is then systematically renamed for clarity, reducing redundancy from overlapping motif characteristics (Table S7). In the volcano plot, each cluster is represented by the point with the highest differential binding score, highlighting the most significant TF binding changes.

### Performance evaluation

The performance of strategies S1 and S2 across different techniques was evaluated by calculating the Area Under the ROC Curve (AUC) using both local and global background modes. During AUC calculation, the footprint score was used as the observed value in S1, while the number of overlapping bases between the footprint and the motif was used as the observed value in S2. The ground truth for bound TF binding sites was defined as motifs reside in the ±50 bp window centered by the corresponding ChIP-seq peak summits. AUC values were calculated for 50 TFs with strong binding signals and compared to assess the predictive performance of each strategy across cFOOT-seq, cFOOT-ATAC-seq, and ATAC-cFOOT-seq.

### Single-Cell Data Analysis

For the single-cell analysis, raw FASTQ files were processed using UMI-tools to tag reads based on the barcode. Following this, the reads were mapped to the reference genome using standard bulk analysis methods. After mapping, the reads were split according to their barcode. The BasalKit^85^ ‘avgmod’ module was used for downstream analysis of each cell, providing information on the conversion rate and depth for each cell. The merged dataset was then used as the reference for downstream analyses, including calculating the Fraction of Reads in Peaks (FRiP) and other statistical metrics.

For each cell, the data were subsequently corrected using the FootTrack. To visualize the TF binding, the corrected conversion rate at motifs identified within ChIP-seq peaks for each TF were plotted. To visualize the TF binding patterns at individual loci, a 5-bp smoothing window was applied to the data. Cells with more than 80% missing values were excluded from further analysis.

### Single-Molecule Data Analysis

Single-molecular footprint analysis provides a more sensitive quantification of TF occupancy at specific loci. For each footprint of interest, reads fully spanning the region were extracted, and the conversion rate of C and G within the footprint were quantified. For the positive strand, footprints were considered unoccupied if the proportion of cytosines converted in the footprint exceeded 10%. Similarly, for the negative strand, footprints were classified as unbound if the proportion of guanines converted exceeded 10%. This approach enables precise assessment of TF binding dynamics at single-molecule resolution.

### ChromHMM analysis

We use the ChromHMM file of HepG2 from ENCODE (accession number: ENCFF808IZE), which contains 18 chromatin states, such as TssA, Tssflank, Tx, TXWk and other regions. For better clustering, we merge TssFlnk, TssFlnkU and TssFlnkD as Tss flank region, EnhG1 and EnhG2 as EnhG region, EnhA1 and EnhA2 as EnhA region, Tx and TxWk as Tx region, for the reason that these regions contain similar histones. Thus, we get 13 chromatin states, and then use K-means clustering method to get the TFOS of transcription factors in different chromatin states.

### Analysis of transcription factor co-occurrence

To enhance the understanding of transcription factor interactions, TF-COMB^54^ was utilized to analyze the co-occurrence among transcription factors. To minimize the inclusion of false positives in the TF pair analysis, the ChIP-seq motifs with a positive Footprint Occupancy Score (FOS) are used as genuine binding sites for generating the robust TF interacting pairs.

To expand the analysis of TF pairs across various conditions, we integrated cFOOT-seq data with FootTrack for de novo prediction of transcription factor binding sites. To emphasize condition-specific TF pairs and reduce the potential for false positives due to motif similarity, we conducted a comparative analysis of TF pairs across different experimental conditions. This method enhances the specificity of the findings by facilitating the identification of TF pairs that are uniquely significant under specific conditions.

### Consensus clustering of TF with varying sensitivity to BRM014 in HepG2

To identify transcription factors with varying sensitivity to BRM014 treatment, we performed consensus clustering on TFs that met stringent filtering criteria. The filtering process involved three steps: (1) TFOS exhibited negative values under any condition across the two replicates are excluded; (2) TFs occurred at least 500 loci overlapping with ATAC ∩ ChIP ∩ SMARC are retained; (3) Outlier filtration, TFOS matrix transform into a matrix of relative changes from Control [Ri = (Ti - T0)/T0]. The DB algorithm (Distance-based outlier detection, *DDoutlier* version 0.1.0) was applied to detect and exclude outliers. Following the above three meticulous steps, we identified 134 qualified TFs for consensus clustering (*ConsensusClusterPlus* version 1.64.0) analysis^88^ (cluster by K-Means, the distance by Euclidean). Notably, the Delta Area revealed an elbow point at k=4, indicating the presence of four distinct clusters with varying response characteristics.

To categorize TFs into BRM014-sensitive and resistant groups independent of their genomic locations. Among the 134 TFs, we segregated the flank change matrix by enhancers and promoters. By repeating the TFOS-based filtering approach, we narrowed down the list to 100 TFs. We combined the enhancer and promoter flank change matrices and performed consensus clustering (K-Means clustering with Euclidean distance).

The resulting four clusters (C1 to C4) exhibited increasing degrees of reduction in chromatin accessibility upon BRM014 treatment. TFs where both enhancer (E) and promoter (P) were classified as C1 were designated as highly resistant TFs. TFs with E and P in C1 or C2 were designated as moderately resistant TFs. TFs displaying E and P in C4 were designated as highly sensitive TFs. Those with E and P intra C3 or C4 were designated as moderately sensitive TFs. All other TFs were categorized as ambiguous.

This nuanced classification scheme provides valuable insights into the differential responsiveness of TFs to BRM014 treatment, highlighting their potential roles in mediating the complex regulatory networks affected by BRM014.

### Assessing TF Regulatory Preferences

To evaluate the regulatory preferences of different TF classes, we first selected all binding sites with a FOS > 0. Next, we calculated the fraction of promoters (±1 kb) from both HepG2-specific and housekeeping genes that overlapped these sites, and then computed the ratio of these fractions.

HepG2-specific genes were obtained from Harmonizome (https://maayanlab.cloud/Harmonizome/), and human housekeeping genes from the Housekeeping and Reference Transcript Atlas (https://housekeeping.unicamp.br).

### Motif Association of Footprints

To assess how many footprints could be linked to known TFs, we first merged adjacent 2-bp windows with footprint scores > 0.05 into contiguous regions of 5–100 bp, designating these as bona fide footprints. Footprints with < 50 % overlap with known motifs in open-chromatin regions were classified as footprints without known motif, while those with ≥ 50 % overlap were classified as footprints with known motif. We then compared the length distributions and footprint scores of these two footprint types to uncover differences in their binding characteristics.

### Data availability

Data related to motifs were sourced from JASPAR^49^ (https://jaspar.elixir.no), while ChIP-seq and ATAC-seq datasets were obtained from Cistrome^89^ (http://cistrome.org/db/#/) and ENCODE (https://www.encodeproject.org/). The specific data sources can be found in the Table S1. Full benchmarking data and updated resources will be available in the peer-reviewed version

### Code Availability

The FootTrack analysis framework and associated scripts used in this study will be made publicly available upon formal publication.

## Notes

### Competing Interest Statement

The authors have declared no competing interest.

