## Supplementary figures for "Genome-Wide Investigation of Transcription Factor Occupancy and Dynamics Using cFOOT-seq"

#### Figure S1

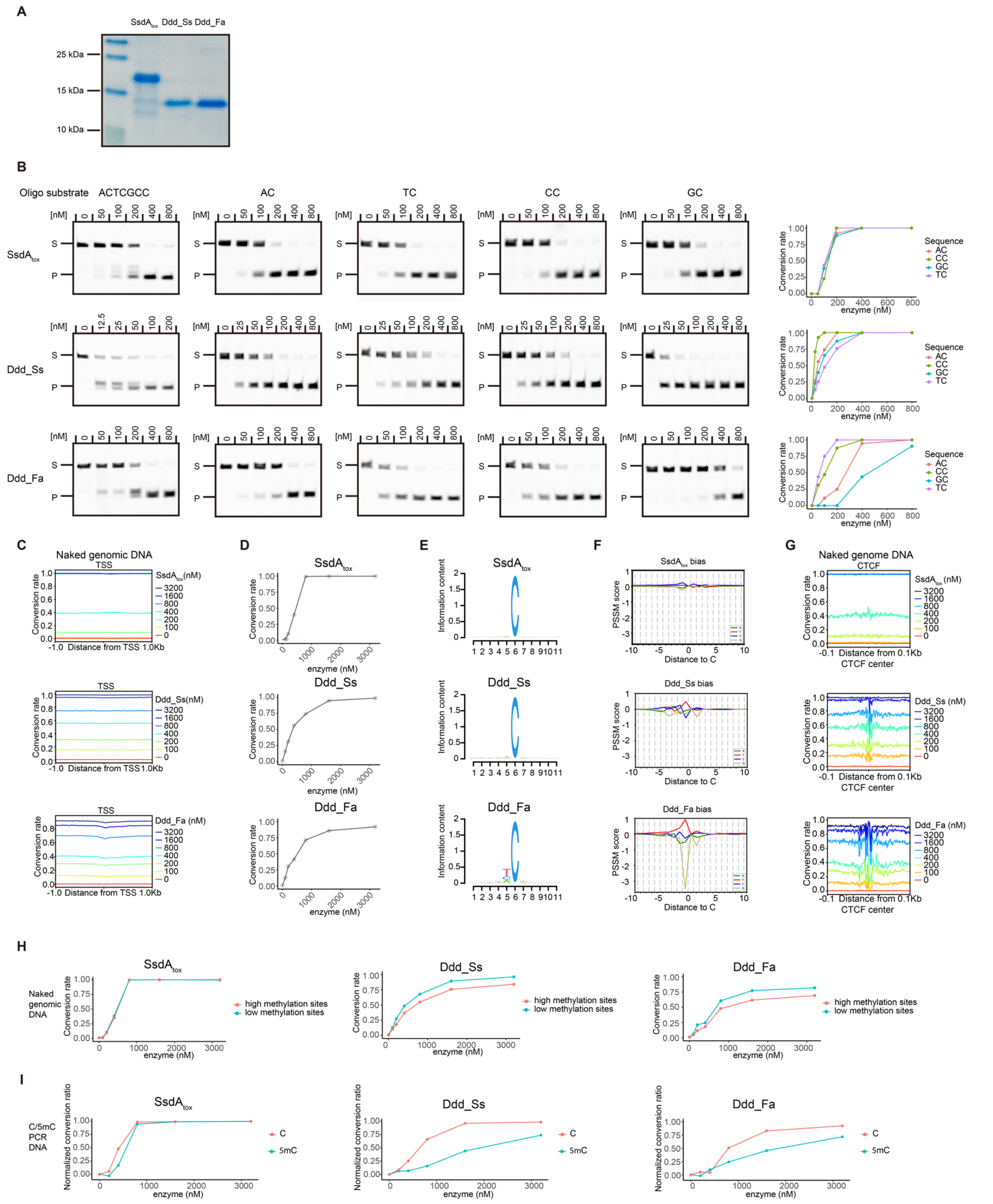


**Figure S1: Deaminase activity and sequence bias of SsdA$_{tox}$, Ddd_Ss, and Ddd_Fa across different substrates, related to Figure 1.**

**A.** Gel image showing purified SsdA$_{tox}$, Ddd_Ss, and Ddd_Fa proteins.

**B.** In vitro cytidine deamination assays using various dsDNA oligonucleotide substrates (ACTCGCC, AC, TC, CC, and GC). Left: Gel images showing the products of deamination. Right: Quantification of deamination efficiency for each substrate, treated with SsdA$_{tox}$ (top), Ddd_Ss (middle), and Ddd_Fa (bottom).

**C.** Average conversion rates around transcription start sites (TSS) in R1 naked genomic DNA treated with increasing concentrations of SsdA$_{tox}$ (top), Ddd_Ss (middle), and Ddd_Fa (bottom).

**D.** Average conversion rates in R1 naked genomic DNA treated with increasing concentrations of SsdA$_{tox}$ (top), Ddd_Ss (middle), and Ddd_Fa (bottom).
**E.** Sequence logos displaying the sequences flanking converted cytosines in R1 naked genomic DNA treated with SsdA$_{tox}$ (top), Ddd_Ss (middle), and Ddd_Fa (bottom).
**F.** Sequence bias for SsdA$_{tox}$ (top), Ddd_Ss (middle), and Ddd_Fa (bottom) on R1 naked DNA before bias correction. The position-specific scoring matrix (PSSM) shows nucleotide preferences from -10 to +10 bp relative to cytosine.
**G.** Average conversion rates around CTCF motifs derived from CTCF ChIP-seq peaks in R1 naked genomic DNA treated with increasing concentrations of SsdA$_{tox}$ (top), Ddd_Ss (middle), and Ddd_Fa (bottom).
**H.** Average conversion rates at high methylation sites (top 50% methylation) versus low methylation sites (bottom 50% methylation) in R1 naked genomic DNA treated with increasing concentrations of SsdA$_{tox}$ (left), Ddd_Ss (middle), and Ddd_Fa (right).
**I.** Conversion rates of C and 5mC PCR DNA substrates treated with increasing concentrations of SsdA$_{tox}$ (left), Ddd_Ss (middle), and Ddd_Fa (right), measured by UPLC-MS/MS

#### Figure S2

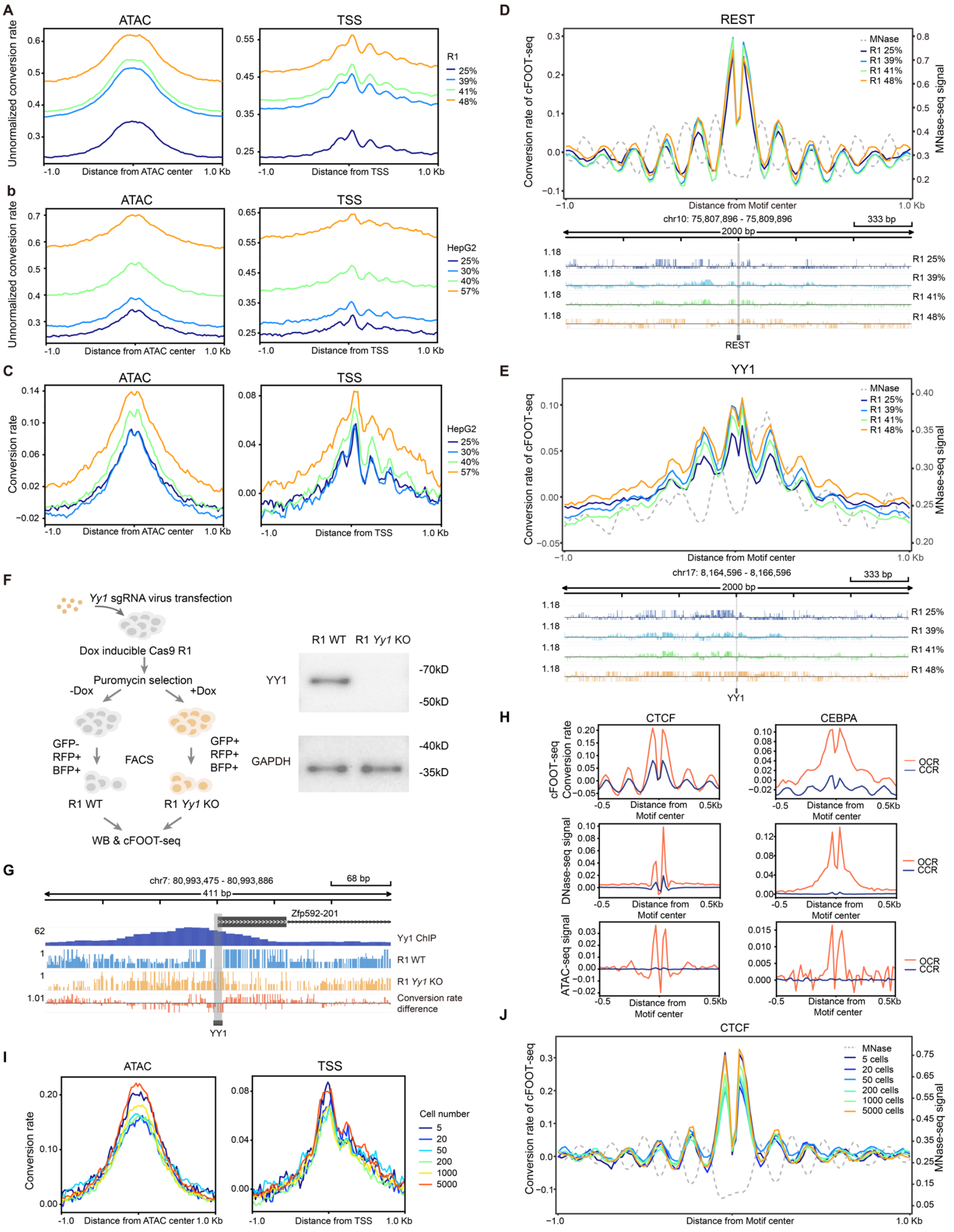


**Figure S2: cFOOT-seq maps chromatin accessibility, nucleosome occupancy, and TF footprint, related to Figure 1.**

**A.** Unnormalized average conversion rates around ATAC-seq peak center (left) and transcription start sites (TSS) (right) in R1 cells with increasing average genomic conversion rates.

**B.** Unnormalized average conversion rates around ATAC-seq peak center (left) and transcription start sites (TSS) (right) in HepG2 cells with increasing average genomic conversion rates.

**C.** Normalized average conversion rates around ATAC-seq peak center (left) and transcription start sites (TSS) (right) in HepG2 cells with increasing average genomic conversion rates.

**D-E.** Average profiles of normalized DNA conversion rates of cFOOT-seq and MNase-seq signal around all REST (E) or YY1 (F) binding sites defined by ChIP-seq aligned to their respective motifs (top), and distribution of DNA conversion rates at a representative genomic region flanking the CTCF or YY1 motif (bottom) in R1 cells. The pattern of nucleosome positioning as indicated by MNase-seq is shown with a grey dashed line. The profiles indicate binding events at motif centers and nucleosome patterns around REST or YY1 binding sites, showing samples with increasing average genomic conversion rates.

**F.** Schematic of *Yy1* KO R1 cell collection through FACS after Dox-induced expression of Cas9 (left). Western blot analysis of YY1 protein levels in cells with or without Dox induction, using antibodies against YY1 and GAPDH (right).

**G.** Distribution of DNA conversion rates around a representative YY1 binding site in R1 WT (33%) and YY1 KO (32%) cells. The YY1 ChIP-seq track (purple) shows the YY1 binding sites, and the position of the YY1 motif is indicated. cFOOT-seq conversion rates for R1 WT (blue) and R1 YY1 KO (orange) cells are shown. The conversion rate difference track (red) represents the difference in conversion rates between R1 WT and R1 YY1 KO cell, reflecting changes in TF binding at the YY1 motif

**H.** Average signal profiles of cFOOT-seq (top), DNase-seq (middle), and ATAC-seq (bottom) in open chromatin regions (OCR) and closed chromatin regions (CCR) around the centers of all CTCF (left) or CEBPA (right) motif center(±0.5kb).

**I.** Normalized DNA conversion rates around ATAC-seq peak center, TSS (±0.1kb) in R1 samples with low input cell numbers (5-5000).

**J.** Average profiles of normalized DNA conversion rates of cFOOT-seq and MNase-seq signal around CTCF binding sites aligned to CTCF motifs in R1 samples with low input cell numbers (5-5000).

#### Figure S3

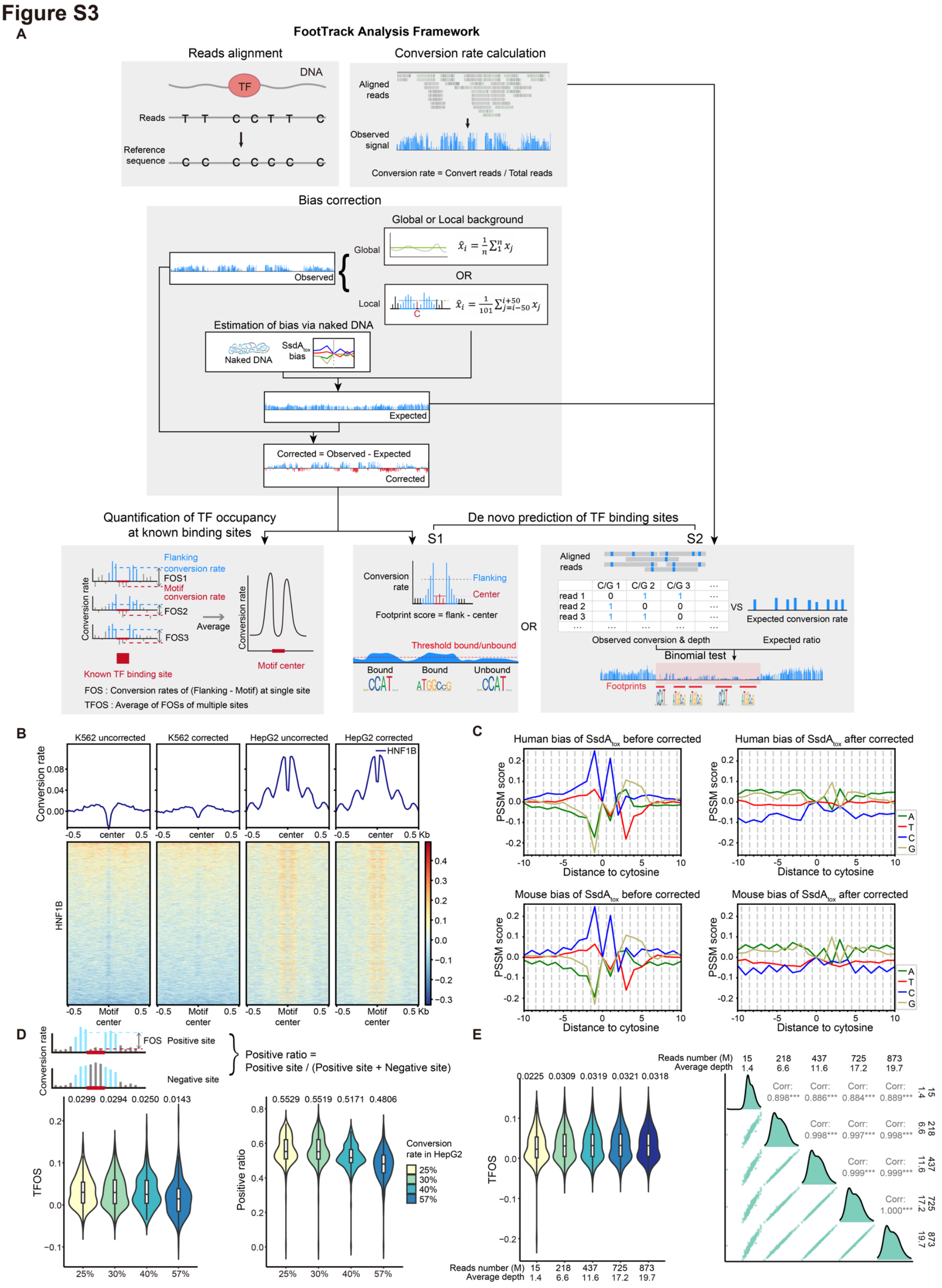


**Fig.S3: FootTrack analysis framework and performance of cFOOT-seq under different conditions, related to Figure 2.**

**A.** Analysis framework for cFOOT-seq processing, including read alignment, C/T conversion calling, bias correction, and downstream analysis. Bias correction is performed using two modes: global background (average conversion rate across the entire genome) and local background (average conversion rate within a ±50 bp window around each base). The corrected conversion rate is calculated by subtracting the expected from the observed conversion rate. The framework supports two primary analyses: (1) Quantification of TF occupancy at known binding sites, (2) De novo prediction of TF binding sites using strategy S1 (calculating footprint scores directly from corrected data) and S2 (predicting footprints using binomial statistical tests)

**B.** Average profiles and heatmaps of normalized DNA conversion rates around HNF1B binding sites in K562 and HepG2 cells, comparing the data before and after bias correction.

**C.** Sequence bias of $SsdA_{tox}$ on naked DNA of human and mouse before and after bias correction. Each panel shows the position-specific scoring matrix (PSSM) scores for nucleotides A, T, C, and G across positions from -10 bp to +10 bp relative to cytosine.

**D.** Schematic illustration of the calculation of the positive ratio of FOS (top), along with violin plots showing TFOS scores (left) and the proportion of sites with positive FOS for each TF (right) under different genomic conversion rates in HepG2 cells using cFOOT-ATAC-seq. The conversion rates are color-coded and labeled at the bottom of each plot. The median values of TFOS and FOS positive ratio are shown above each violin plot.

**E.** Violin plots (left) and correlation plots (right) showing TFOS at various sequencing depths in HepG2 cells. The median TFOS values for all tested TFs in HepG2 is indicated above each violin plot. The total sequencing amount and average reads depth for each condition is listed for the corresponding plots.

#### Figure S4

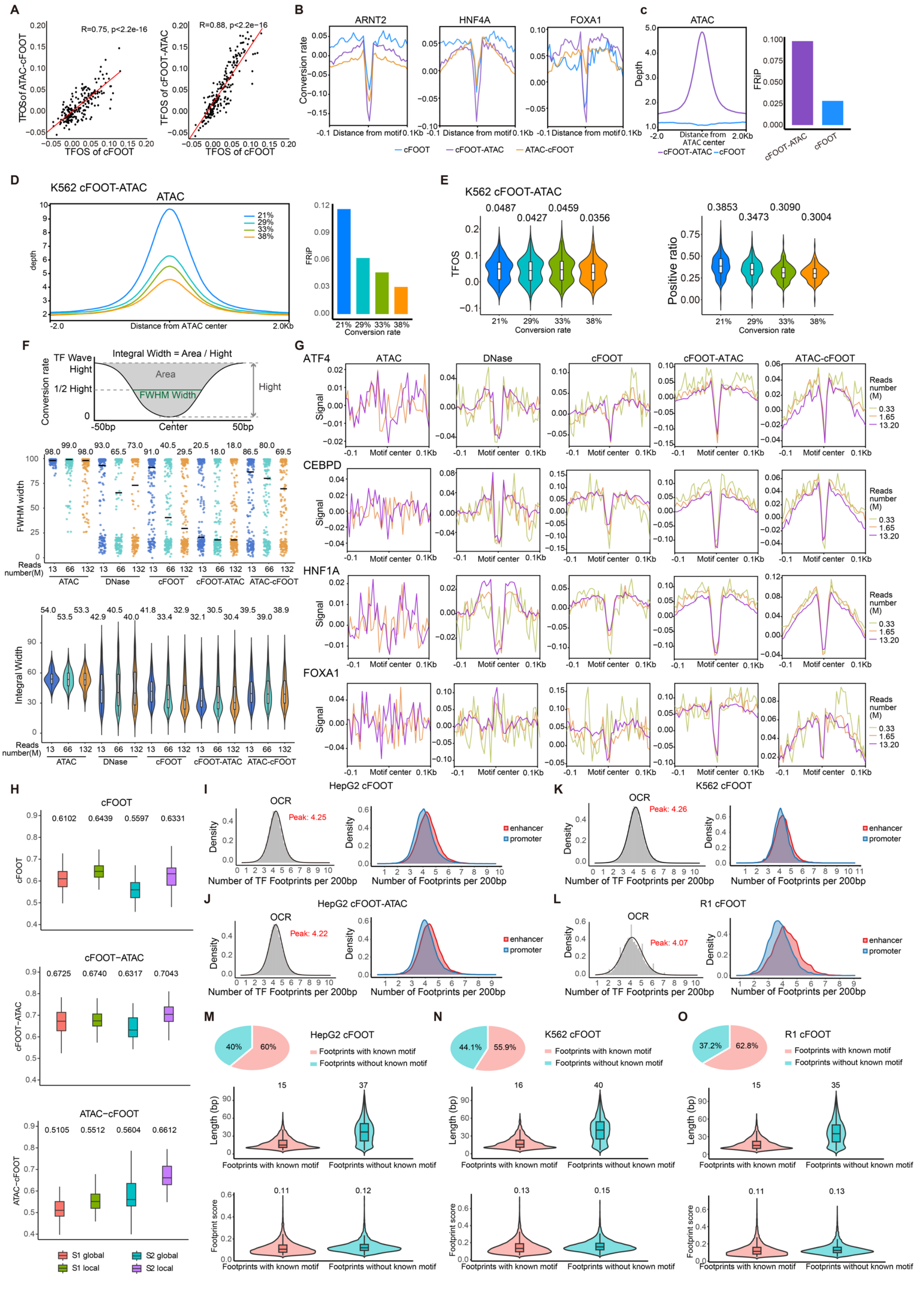


**Fig.S4 cFOOT-seq combined with ATAC-seq provide high-resolution and sensitive view of genomic TF occupancy, related to Figure 2.**

**A.** Scatter plot showing the correlation between TFOS differences of cFOOT-seq (30%) and ATAC-cFOOT-seq (33%) (R=0.75, p < 2.2e-16), cFOOT-seq (30%) and cFOOT-ATAC-seq (36%) (R=0.88, p < 2.2e-16) for each transcription factor (TFOS -0.05~0.20) in HepG2 cells.

**B.** Normalized average conversion rates around ±0.1 kb of ARNT2 (left), HNF4A (middle), and FOXA1 (right) motifs in HepG2 cells, comparing conversion rates from cFOOT-seq, cFOOT-ATAC-seq, and ATAC-cFOOT-seq methods.

**C.** Comparison of cFOOT-ATAC-seq and cFOOT-seq methods in HepG2 cells showing read depth around the ATAC-seq peak center (left) and FRiP values (right). Data for all conditions were obtained by extracting 10 million reads.

**D.** Comparison of cFOOT-ATAC-seq (29%) depth at ATAC-seq peak center with different conversion rates in K562 cells (left), and the corresponding FRiP values (right). Data for all conditions were obtained by extracting 10 million reads.

**E.** Violin plots showing TFOS scores (left) and the proportion of sites with positive FOS for each TF (right) under different genomic conversion rates in K562 cells using cFOOT-ATAC-seq. The conversion rates are color-coded and labeled at the bottom of each plot. The median values of TFOS and positive FOS ratio for all tested TFs are indicated above each violin plot.

**F.** Schematic representation (top) of the calculation for integral width and full-width half-maximum (FWHM) width of TF footprints. The scatter plot (middle) and violin plot (bottom) show the distribution of FWHM width and integral width, respectively, illustrating the resolution of ATAC-seq, DNase-seq, cFOOT-seq, cFOOT-ATAC-seq, and ATAC-cFOOT-seq methods in detecting 204 HepG2 TF footprints across different sequencing depths. The median values of FWHM width and integral width of each tested method are shown above the graphs.

**G.** Normalized average conversion rates around ±0.1 kb of ATF4, CEBPD, HNF1A, FOXA1 motif center in HepG2 cells. The plot compares the ability of ATAC-seq, DNase-seq, cFOOT-seq, cFOOT-ATAC-seq, and ATAC-cFOOT-seq methods to detect these four TF footprints under different reads number (0.33 M, 1.65 M, 13.20 M). The signal of ATAC-seq with 0.33 M is not available due to low reads number

**H.** Area under the ROC curve (AUC) for both strategies (S1 and S2) using local and global background modes. Results are shown for cFOOT-seq (left), cFOOT-ATAC-seq (middle), and ATAC-cFOOT-seq (right) of HepG2.

**I-L.** Density distributions of de novo predicted footprints (per 200 bp) by FootTrack in open chromatin regions (OCRs, defined by ATAC-seq), with comparisons between promoter-associated and enhancer-associated OCRs in HepG2 cFOOT-seq (I), HepG2 cFOOT-ATAC-seq (J), K562 cFOOT-seq (K), and R1 cFOOT-seq (L).

**M-O.** Pie charts show the proportions of footprints in OCRs with or without known motifs. Violin plots display the distributions of footprint lengths (middle) and footprint scores (bottom) for each category in HepG2 cFOOT-seq (M), K562 cFOOT-seq (N), and R1 cFOOT-seq (O).

#### Figure S5

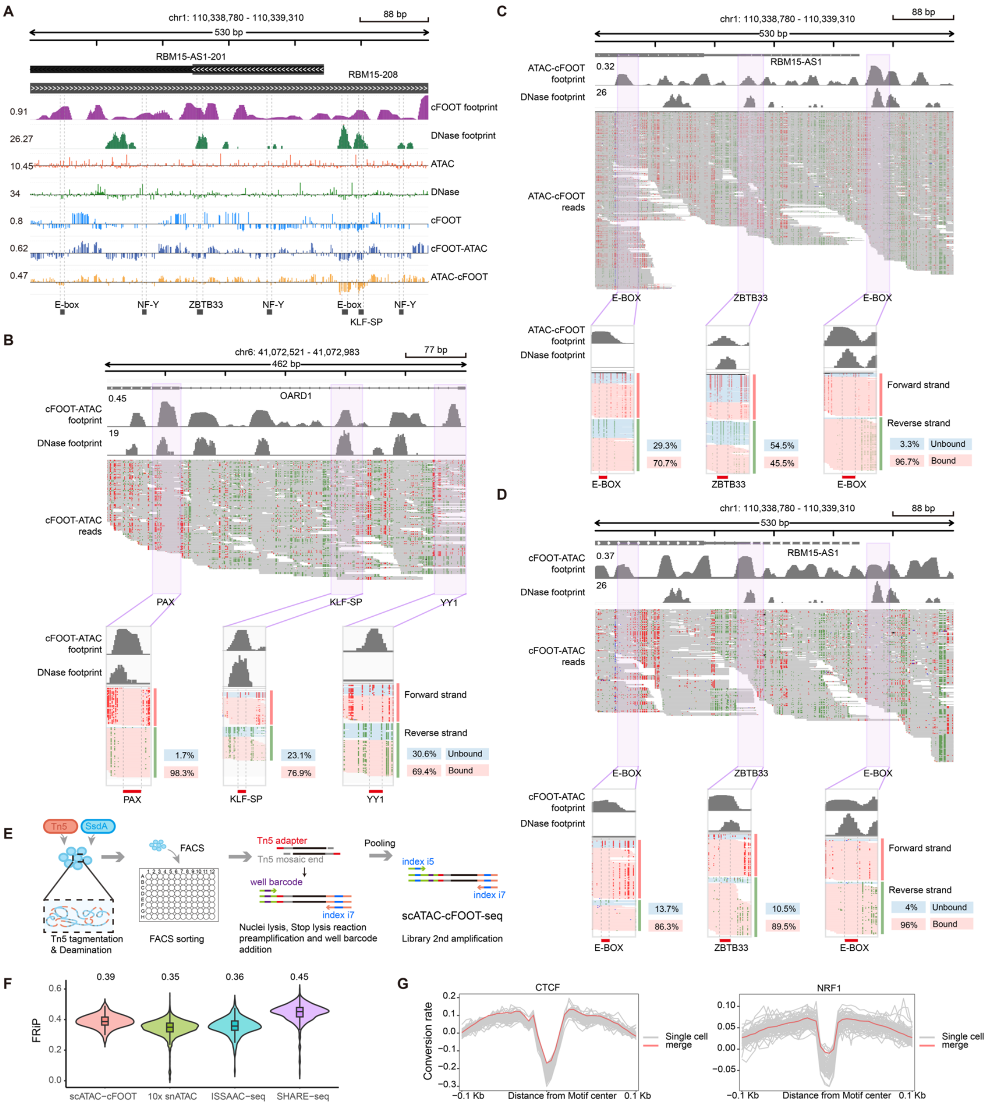


**Figure S5: Single-molecule and single-cell analysis of cFOOT-seq combinational methods, related to Figure 2.**

**A.** IGV browser graphics showing profiles of cFOOT-seq, cFOOT-ATAC-seq, ATAC-cFOOT-seq, DNase-seq, and ATAC-seq at a representative site (chr1: 110,338,780–110,339,310) in HepG2 cells. The "cFOOT footprint" track shows footprint scores derived from cFOOT-seq data, while the "DNase footprint" track shows footprint probabilities from DNase-seq data. The five tracks below display the distribution of corrected cutting events for ATAC-seq and DNase-seq, along with the corrected conversion rates for cFOOT-seq, cFOOT-ATAC-seq, and ATAC-cFOOT-seq. The black labels at the bottom highlight motifs identified in ChIP-seq peaks, corresponding to TF binding sites

**B.** IGV browser visualization showing the single-molecule profile of cFOOT-ATAC-seq at a representative site (chr6: 41,072,521–41,072,983) in HepG2 cells. The "cFOOT-ATAC footprint" track represents footprint scores calculated from cFOOT-ATAC-seq data, while the "DNase-seq footprint" track shows footprint probabilities derived from DNase-seq data. The " cFOOT-ATAC reads" track displays individual mapped reads in this region. Both forward and reverse strands are shown for the three TF motifs (PAX, KLF-SP, and YY1), and categorized into bound and unbound states based on conversion rates at each motif

**C-D.** IGV browser visualization showing the single-molecule profile of of ATAC-cFOOT-seq data (C) and cFOOT-ATAC data (D) at a representative site (chr1: 110,338,780–110,339,310) in HepG2 cells. The " cFOOT-ATAC footprint" track represents footprint scores calculated from cFOOT-ATAC-seq data, while the "DNase footprint" track shows footprint probabilities derived from DNase-seq data. The " cFOOT-ATAC reads" track displays individual mapped reads in this region. Both forward and reverse strands are shown for the three TF motifs (two E-boxes, and ZBTB33), and categorized into bound and unbound states based on conversion rates at each motif

**E.** Schematic of the experimental workflow for scATAC-cFOOT-seq, including Tn5 fragmentation, deamination, FAC sorting, barcoding, and pooling for library construction.

**F.** Violin plots showing the distribution of FRiP (fraction of reads in peaks) for single cells in scATAC-cFOOT-seq, 10x snATAC-seq, ISSAAC-seq and SHARE-seq of K562.

**G.** Plot showing the conversion rate at aggregated CTCF (left) and NRF1 (right) binding motifs in individual single cells (gray lines) and the merged data (red line) of K562.

#### Figure S6

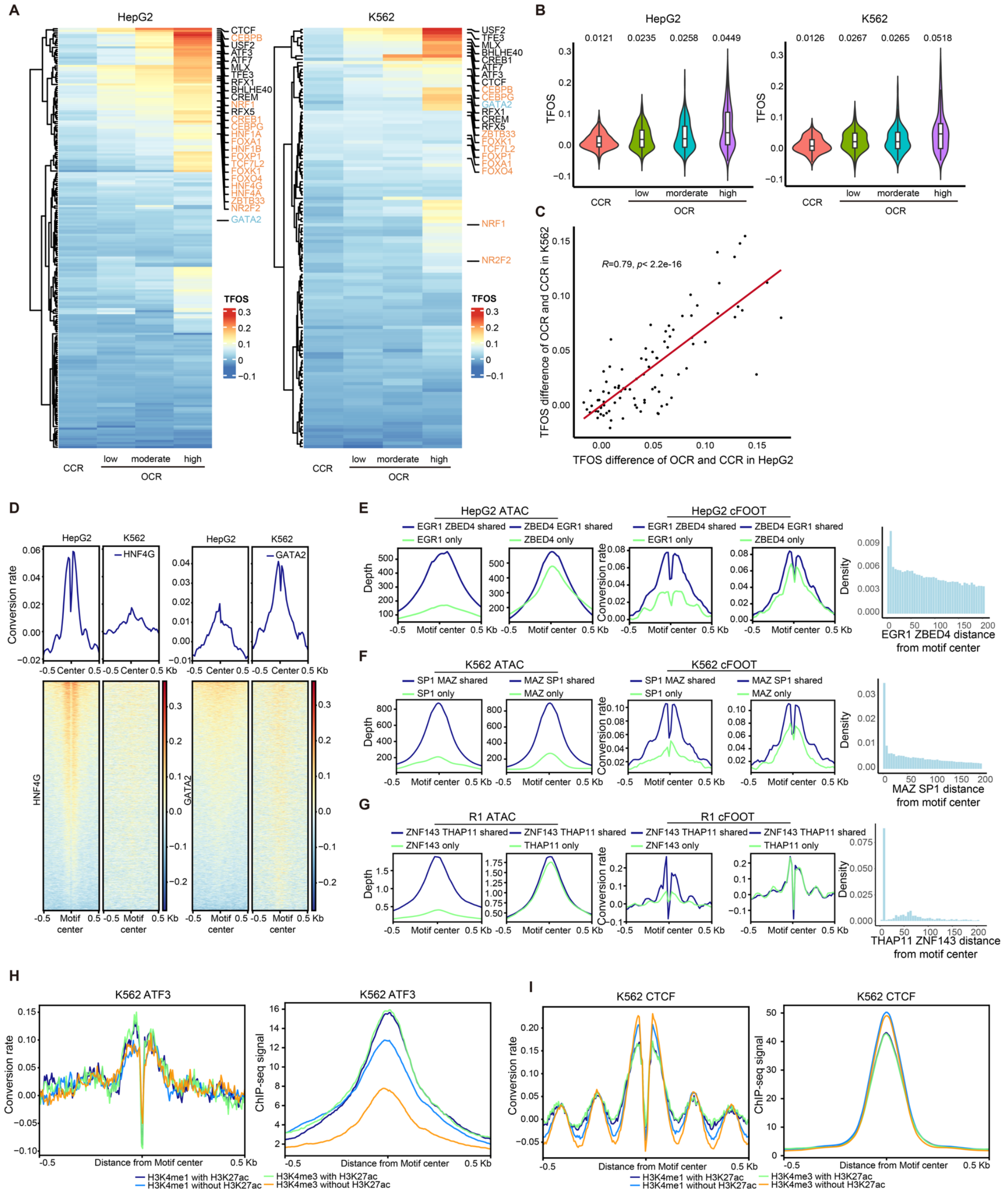

**Figure S6: cFOOT-seq quantitatively assess the impact of chromatin accessibility, histone modifications, and cofactors on TF occupancy, related to Figure 3.**

**A.** Heatmaps showing TFOS across different chromatin accessibility regions in HepG2 (left, 30%) and K562 (right, 30%) cells. Regions are categorized into closed chromatin regions (CCR) and low, moderate, and high open chromatin regions (OCR). TFs highlighted in orange indicate those with higher TFOS in OCR of HepG2, while TFs highlighted in blue indicate those with higher TFOS in OCR of K562. The color scale represents TFOS depth, with higher values in red and lower values in blue.

**B.** Violin plots illustrating the distribution of TFOS scores in different chromatin accessibility regions (CCR, low, moderate, high OCR) for HepG2 (left) and K562 (right) cells. The median TFOS values for each regioon are indicated above each violin plot.

**C.** Scatter plot showing the correlation between TFOS differences of OCR and CCR for each transcription factor in HepG2 and K562 cells. (n=89, R=0.79, p < 2.2e-16).

**D.** Average profiles and heatmaps displaying corrected DNA conversion rates around the motif centers for HNF4G and GATA2 binding sites defined by ChIP-seq in HepG2 and K562 cells. The differences in conversion patterns between HepG2 and K562 cells indicate cell type-specific flanking chromatin accessibility and TF occupancy around the motif centers for HNF4G and GATA2

**E.** Average profiles showing ATAC-seq reads depth (left) or corrected DNA conversion rates (middle) around ZBED4 and EGR1 binding sites in HepG2 cells, and the density distribution of distances between EGR1 and ZBED4 motif centers (right).

**F**. Average profiles showing ATAC-seq reads depth (left) or corrected DNA conversion rates (middle) around MAZ and SP1 binding sites in K562 cells, and the density distribution of distances between MAZ and SP1 motif centers (right).

**G**. Average profiles showing ATAC-seq reads depth (left) or corrected DNA conversion rates (middle) around ZNF143 and THAP11 binding sites in R1 cells, and the density distribution of distances between ZNF143 and THAP11 motif centers (right).

**H-I.** Average profiles showing corrected DNA conversion rate (left) and ChIP-seq signal (right) around ATF3 (**H**) and CTCF (**I**) binding sites in K562 cells. Binding sites were defined by ChIP-seq of each TF. Both cFOOT-seq and ChIP-seq signals are shown separately for regions marked by different combinations of histone modifications (H3K4me1$^{+}$ H3K27ac$^{-}$, H3K4me1$^{+}$ H3K27ac$^{+}$, H3K4me3$^{+}$ H3K27ac$^{-}$, and H3K4me3$^{+}$ H3K27ac$^{+}$) of K562 cells.

**Figure S7**

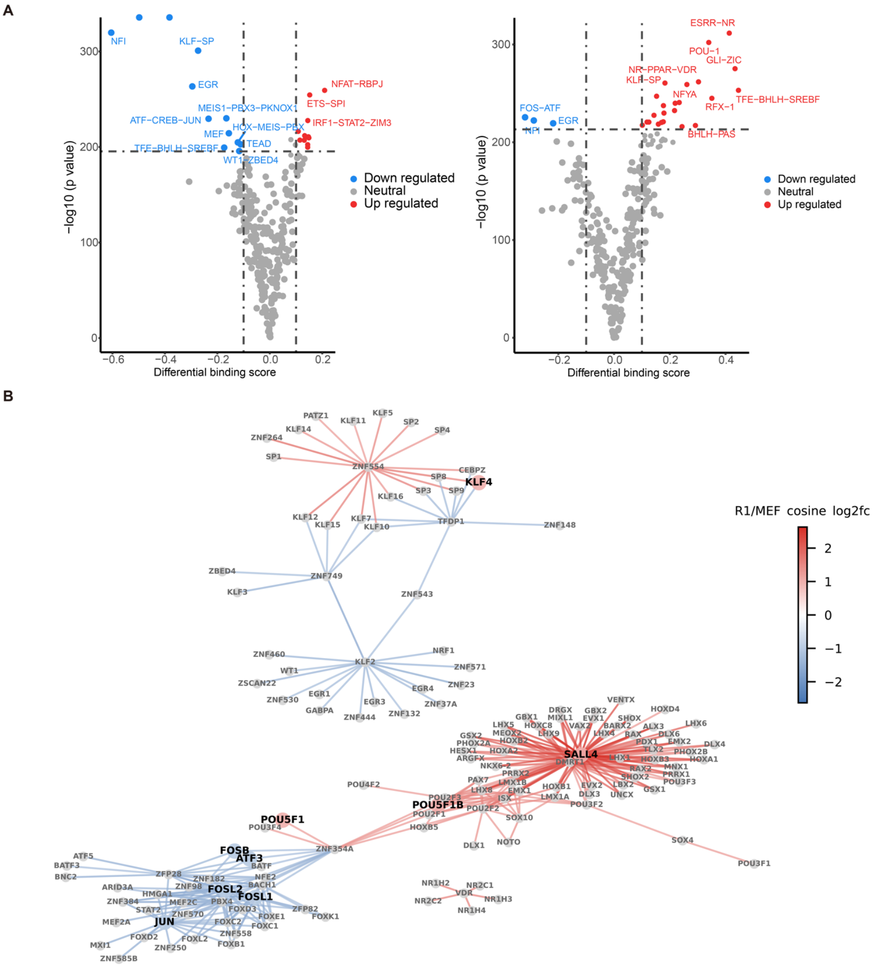


### Figure S7: Detailed analysis of TF binding differences in MEF and R1, related to Figure 4.

**A**. Volcano plots showing FootTrack-predicted TF clusters with differential TF binding scores between MEF and R1 cells in MEF open regions (left) and R1 open regions (right). TFs with significantly upregulated footprint scores in R1 cells are highlighted in red, while those significantly downregulated in R1 are highlighted in blue.

**B**. Transcription factor co-occurrence network changes Between MEF and R1 Cells. This network diagram illustrates the co-occurring transcription factor (TF) pairs with edges colored based on the cosine log2 fold change (log2fc) between R1 and MEF cells. Nodes representing TFs specific to R1 are highlighted in red, while those specific to MEF are highlighted in blue, indicating differential TF associations unique to each cell type.

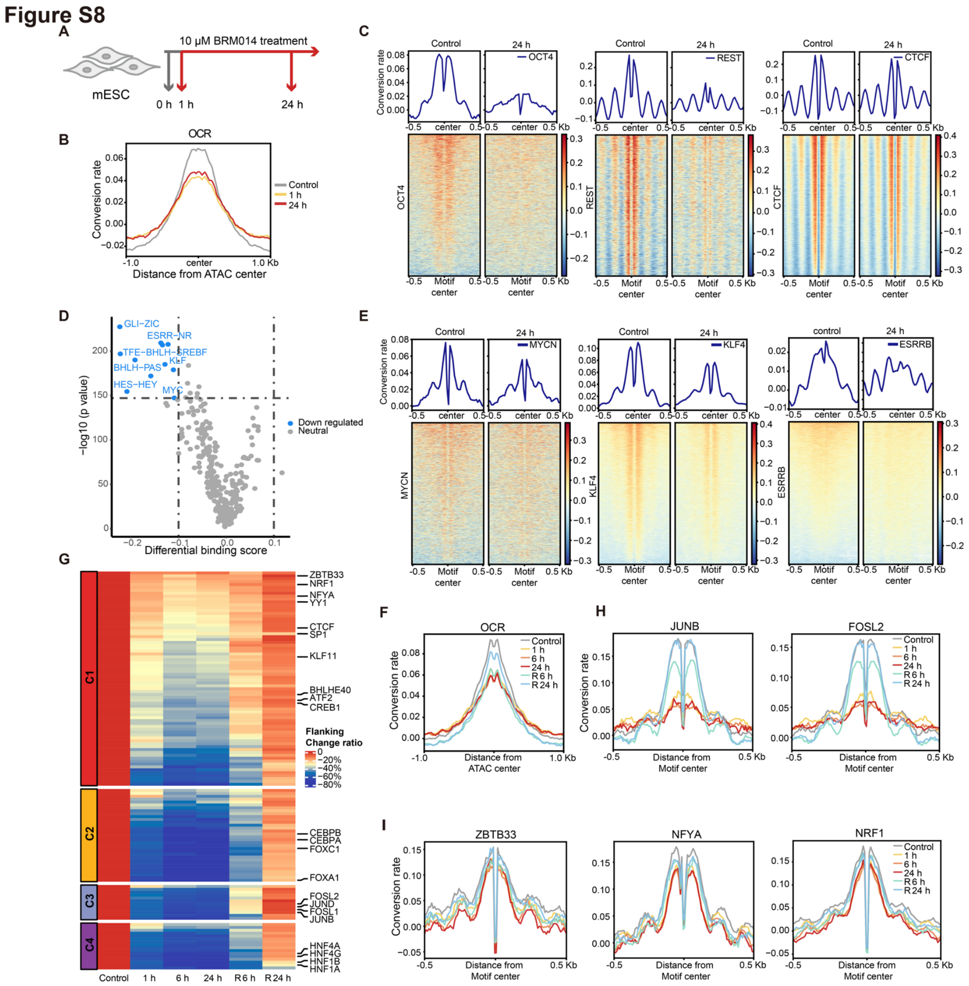


**Figure S8: cFOOT-seq depicts dynamics of nucleosome organization and TF occupancy in response to inhibition of SWI/SNF, related to Figure 5.**

**A.** Schematic representation of BRM014 treatments on mESCs (OG2 cells).

**B.** Average profile showing normalized DNA conversion rates around OCR defined by ATAC-seq in mESCs (OG2 cells) treated with BRM014 at 0 h (control, 39%), 1 h (40%), and 24 h (38%).

**C.** Average profiles and heatmaps showing DNA conversion rates around OCT4, REST, and CTCF binding sites under control conditions and after 24 h BRM014 treatment. OCT4 and REST sites show decreased chromatin accessibility and altered TF footprints, while CTCF sites exhibit minimal change.

**D.** Volcano plots showing FootTrack-predicted TF clusters with differential TF binding scores between control and 24 h BRM014-treated OG2 cells at open regions. TFs with downregulated scores are shown in blue, indicating decreased binding activity.

**E.** Average profiles and heatmaps of conversion rates around MYCN and KLF4 binding sites defined by ChIP-seq in OG2 cells treated with BRM014 for 0 h (control) and 24 h, aligned to motif centers and ranked by average conversion rates.

**F.** Average profiles of DNA conversion rates around OCR illustrating changes in chromatin accessibility following different durations of BRM014 treatment and recovery in HepG2 cells.

**G.** Heatmap showing the dynamic changes in the flank accessibility of 4 categories of TFs responding to BRM014 as defined by TFOS change ratio, with the left label marked the consensus clusters derived from TFOS (as shown in 5C), and the colors represent the proportion of flank accessibility change relative to the 0 h untreated baseline.

**H-I.** Average profiles of DNA conversion rates around TF binding sites of JUNB and FOSL2 (H), ZBTB33, NFYA and NRF1 (I) under control, 1 h, 6 h, and 24 h BRM014 treatments, as well as 6 h and 24 h recovery after 24 h BRM014 treatment. The profiles show changes in chromatin accessibility and TF footprints across different treatment conditions.

**Figure S9**

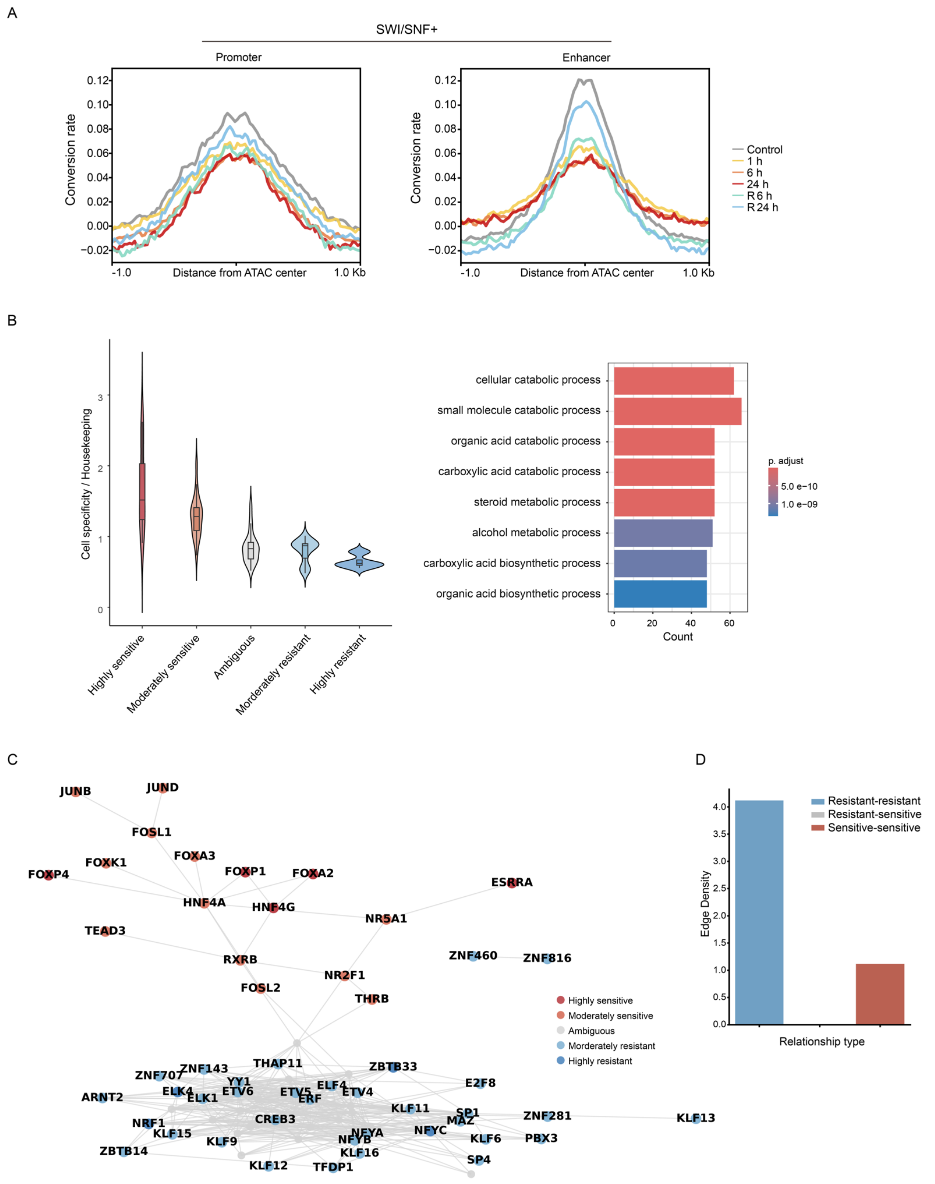


**Figure S9: Definition of TF dependency on SWI/SNF reveals spatial organization rule of TFs, related to Figure 6.**

**A.** Average profiles of DNA conversion rates in promoter (left) and enhancer (right) around SWI/SNF$^+$ OCR regions, illustrating changes in chromatin accessibility following different durations of BRM014 treatment and recovery.

**B.** Violin plot showing, for each TF class, the ratio between the proportion of TF binding at cell-specific gene promoters and that at housekeeping gene promoters (left). GO enrichment of HepG2-specific genes (right).

**C.** This network diagram displays the co-occurrence of transcription factor (TF) pairs within the promoter regions of HepG2 cells. Each node represents a different TF, with the color coding indicating varying levels of sensitivity or resistance. Edges between nodes indicate potential co-occurrence relationships among the TFs

**D.** The bar graph displays the edge density in transcription factor co-occurrence network within the promoter regions of HepG2 cells among transcription factor pairs, categorized by their sensitivity to BRM014. The categories include interactions between TF pairs that are both resistant (blue bar), one resistant and one sensitive (grey bar), and both sensitive (red bar).
